# Calneuron 1 reveals the pivotal roles in schizophrenia via perturbing human forebrain development and causing hallucination-like behavior in mice

**DOI:** 10.1101/2024.04.16.589839

**Authors:** Hui-Juan Li, Xiao Yu, Xi Liu, Jinhong Xu, Jinlong Chen, Tianlin Cheng, Sangmi Chung, Yousheng Shu, Zhicheng Shao

**Author notes:** These authors contributed equally (Hui-Juan Li, Xiao Yu, Xi Liu).

## Abstract

Schizophrenia is a highly heritable neurodevelopmental disorder with unknown genetic pathogenic mechanisms. Here, we selected 11 schizophrenia risk genes and generated single-gene-knockout-precise-dorsal/ventral-forebrain-organoids (SKOPOS) via CRISPR-Cas9 system. 90 bulk and 249,430 single-cell RNA-sequencing of SKOPOS revealed that knockout of 11 risk genes lead to different levels of deficits in dorsal/ventral forebrain organoids. Among them, calneuron 1 (*CALN1*) acts as a pivotal pathogenic gene of schizophrenia via severe disruption of gene expression network, interaction with about 32% (34/106) known schizophrenia risk genes, delayed maturation and impaired spontaneous neural circuit in human developing forebrain. Furtherly, including the spontaneous abrupt burst spiking in cortical neurons and the defects of spatial memory, cognition and social ability, *Caln1* KO mice surprisingly displayed spontaneous startle behavior and head-twitch response correlated with hallucination-like behavior, which could be inhibited by antipsychotic drug SEP-363856. In summary, *CALN1* is identified as a pivotal pathogenic gene of schizophrenia in forebrain development.

## Introduction

Schizophrenia (SCZ) is a highly heritable neurodevelopmental disorder including impairments of cognition, perception and motivation that usually affects about 1% of the world’s population^1–3^, and previous studies have revealed that the heritability of this illness was estimated to be about 80%^4,5^. A recent study has prioritized 106 protein-coding genes as risk genes by fine-mapping and functional genomic analysis based on genome-wide association study (GWAS) of 76,755 schizophrenia patients and 243,649 control individuals, opening an avenue to understand schizophrenia pathogenic mechanisms^6^. Among these genes, several genes have been reported to be well related to neurological function, for example *GRIN2A*^7,8^, but the roles of most risk genes in the nervous system are unknown. Deciphering how these risk genes disturb brain function and finally how to contribute to SCZ pathogenesis is still a huge challenging task.

Human forebrain plays a central role in complex cognitive activities, sensory function and motor activities, and has been showed structural and functional abnormalities in patients diagnosed with schizophrenia^9–12^. To overcome the challenges of the accessibility of developing human brain tissue, brain organoids derived from human pluripotent stem cells have great potential to investigate human brain development and the pathology of neuropsychiatric disease^13–15^. Using cerebral organoids generated from patient-derived induced pluripotent stem cells (iPSCs), Kathuria and Notaras *et al.* explored the neuropathology of schizophrenia and revealed dysregulated gene expression patterns in cases^16,17^. Besides, Sebastian *et al.* revealed abnormal developmental trajectories and transcriptomic signatures in forebrain organoids derived from schizophrenia patients with *NRXN1* deletions ^18^. These studies further supported that the disturbances in early brain development contribute to the pathogenesis of schizophrenia. However, it is difficult to quantify the contribution to schizophrenia of every genetic variant or risk gene in complex genetic structures of patients ^6,19^, and so we think that the exploration of individually candidate gene role in the early-stage human forebrain development will contribute to understanding of their pathogenic mechanisms.

Herein, through CRISPR-Cas9, we generated single-gene-knockout-precise-dorsal/ventral-forebrain-organoids (SKOPOS) and displayed the roles of 11 risk genes in region-specific human forebrain organoids (DFOs and VFOs) and uncovered the regulatory network of polygenes of SCZ. Among them, *CALN1* may act as a pivotal pathogenic gene of schizophrenia via severe disruption of gene expression network in human developing forebrain, interaction with about 32% (34/106) known schizophrenia risk genes, delayed maturation and impaired spontaneous neural circuit. Importantly, including the spontaneous abrupt burst spiking in cortical neurons and the defects of spatial memory, cognition, social ability and pre-pulse inhibition, KO *Caln1* mice surprisingly displayed spontaneous startle behavior and head-twitch response, which has been reported highly related with hallucination in human^20–25^, as well as can be blocked by antipsychotic drug SEP-363856.

## Results

### Knockout of schizophrenia risk genes in human embryonic stem cell lines

To discover the roles of 106 schizophrenia risk genes in neurodevelopment^6^, firstly, we knocked out pilot 11 genes (*i.e.*, 5 FINEMAP genes and 6 SMR genes) individually in human embryonic stem cells (hESCs) using CRISPR-Cas9 system **(Figure S1a)**. Specifically, these 11 genes include voltage-gated channel *CLCN3*, calcium-binding protein *CALN1*, hormone receptor *CRHR1*, glutamate receptor *GRM1*, glycoprotein *GPM6A*, microtubule protein *MAPT*, splicing factor *SF3B1*, RNA-binding protein *THOC7* and three metabolic enzymes *PLCH2*, *PDE4B* and *PCCB* **(Figure S1a)**. According to the location on the genome, they span ten different genome loci which are genome-wide significantly associated with schizophrenia (P<5×10^−8^) **(Figure S1b-k)**. Based on the BrainSpan database, we found that these 11 risk genes were expressed in both human developmental and adult dorsolateral and ventrolateral prefrontal cortex (**Figure S2a**). After infection of lentivirus including sgRNA, 115 hESC lines were isolated and expanded to individually target these 11 genes **(Figure 1a, b)**. By the verification of Sanger sequencing, these hECS lines include 13 homozygous mutations, 65 heterozygous mutations and 37 wild types (WT) **(Figure 1b)**. The hESC lines with homozygous mutation are distributed across 8 of 11 risk genes **(Figure 1b)**. Next, except negative (normal hESC H9) and positive control (hESC H9 carrying sgRNA of non-target human), we selected one knock-out hESC line per gene including *CLCN3*^−/−^, *CALN1*^−/−^, *CRHR1*^−/−^, *GRM1*^−/−^, *GPM6A*^−/−^, *MAPT*^−/−^, *PLCH2*^−/−^, *PDE4B*^−/−^, *PCCB*^+/−^, *SF3B1*^+/−^ and *THOC7*^+/−^ for downstream experiments in this study **(Figure S3a)**. For each selected KO hESC line, we performed off-targeting assessment using CRISPRoff^26^ and the Sanger sequence furtherly verified that there were no off-target in the top ten predicted genome sites (**Table S1**). And then we also confirmed that the pluripotency of all knock-out hESC lines are maintained, with highly expressing pluripotent markers OCT4 and SSEA4 **(Figure S3b)**.

**Figure 1.**
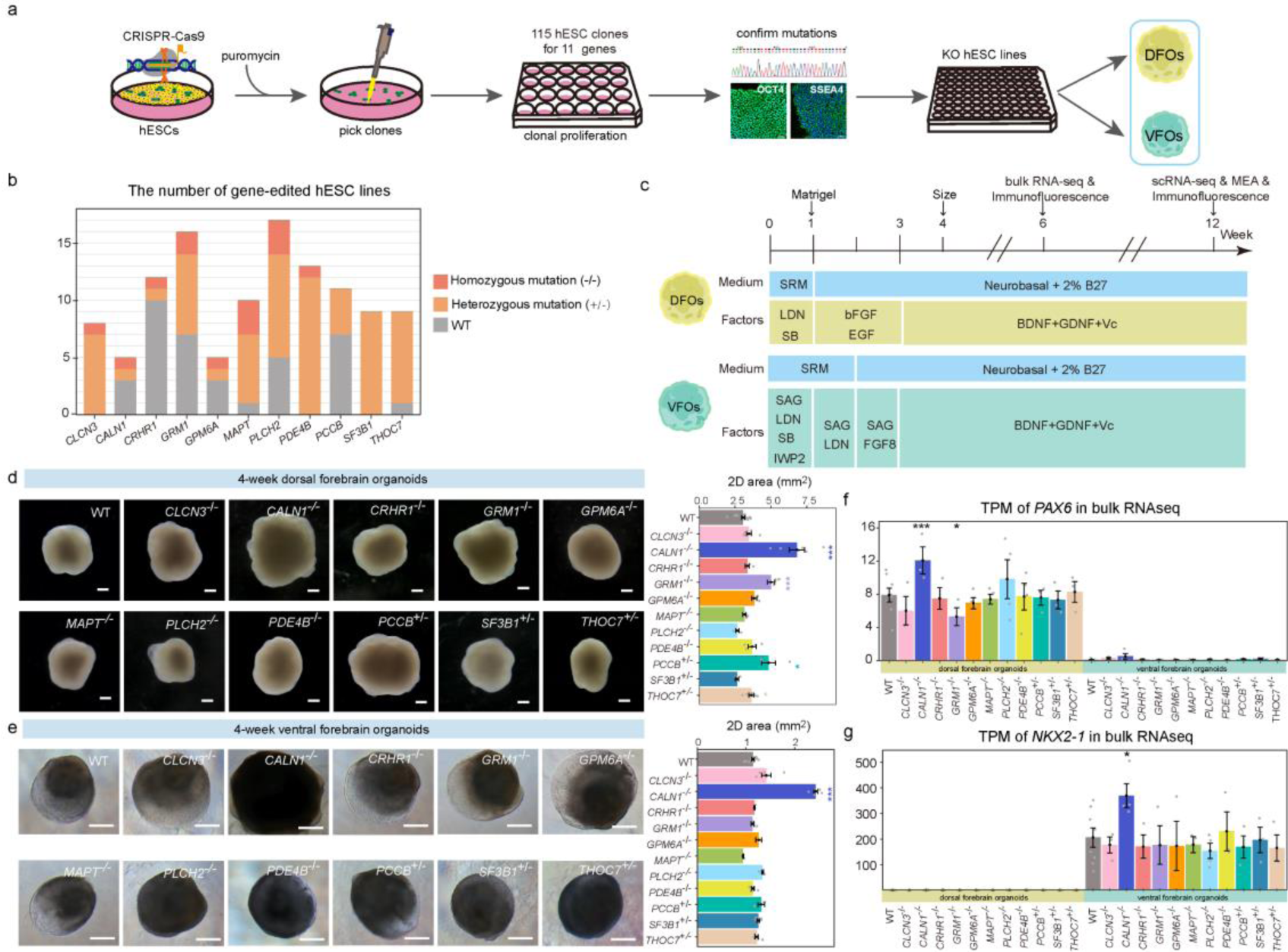
Construction of knock-out hESC lines and dorsal and ventral forebrain organoids. **a**, Schematic illustration for construction of KO cell lines in this study. hESC, human embryonic stem cells. DFOs, dorsal forebrain organoids. VFOs, ventral forebrain organoids. **b**, The number of gene-edited hESC lines verified by Sanger sequence. WT, wild type. **c,** The schematics of generating DFOs and VFOs. LDN, LDN193189. SB, SB431542.Vc, Vitamin C. **d, e,** The size of DFOs (d) and VFOs (e) at 4 weeks. **f**, **g**, The normalized expression values of *PAX6* (**f**) and *NKX2-1* (**g**) of dorsal and ventral forebrain organoids by bulk RNA-seq at 6 weeks. The Wald test was used to differentially expressed analysis. TPM, transcripts per million. Data are presented as mean ± SE. ***, p-value < 0.001. *, p-value < 0.05.

### Generation of dorsal and ventral forebrain organoids as 3D models of schizophrenia risk genes

The expression of genes associated schizophrenia risk variants was enriched in both excitatory neurons and inhibitory neurons^6^, which were respectively generated from the dorsal and ventral telencephalon during human early developmental brain^27^. To fully investigate how schizophrenia risk genes affect early forebrain development in excitatory and inhibitory system, we individually generated DFOs and VFOs induced from 11 KO hESC lines and two wild-type hESC lines **(Figure 1c)**. To reduce the variation of pluripotency, each KO hESC line was used in this experiment between at early passage 3-10. We quantified the size of DFOs/VFOs of each gene at 4 weeks, and observed that both DFOs and VFOs with *CALN1^−/−^*are significantly larger than wild-type organoids at 4 weeks **(Figure 1d, e)**. Besides, DFOs with knockout of *GRM1*, *PDE4B* and *PCCB* are also larger than wild type (**Figure 1d, e**). Using bulk RNA-seq, we found that all DFOs have high expression of *PAX6*, the dorsal forebrain marker, and almost no expression of *NKX2-1*, a ventral forebrain marker, while *NKX2-1* was expressed in every VFO, validating that DFOs/VFOs of each KO gene were induced correctly **(Figure 1f, g).** Bulk RNA-seq or qPCR showed the RNA levels of the target genes in KO 6-week-old organoids, are significantly reduced (**Figure S4a**), and we furtherly confirmed that CALN1 loss at protein level in *CALN1^−/−^* DFOs and VFOs (**Figure S4b**). At the same time, we also observed no expression of Cas9 in 6-week-old organoids, which eliminates the possibility of off-target edits in the stage of organoid differentiation **(Figure S4c)**. To further characterize DFOs and VFOs, we compared the transcriptome of forebrain organoids with human fetal brains **(Figure S4d)**. Firstly, we performed weight gene co-expression network analysis (WGCNA) of 47 RNA-seq datasets from DFOs and 43 RNA-seq datasets from VFOs at 6 weeks. A total of 22 and 19 gene expression modules (except for the gray module) were identified by WGCNA in DFOs and VFOs, respectively **(Figure S4e, f)**. And then from BrainSpan database, we separately downloaded 17 and 16 gene expression profiles of fetal dorsolateral and ventrolateral prefrontal cortex (DLPFC/VLPFC). The Z-summary statistics displayed that all 22 expression modules in DFOs and 16 out of 19 expression modules in VFOs are well-preserved in fetal DLPFC and VLPFC (Z-score > 1.96) **(Figure S4g, h)**. Taken together, these data indicated that DFOs and VFOs with KO schizophrenia risk genes were successfully constructed, and showed similar gene co-expression networks with human fetal brains, which are suitable for modeling and discovering the deficits in the human developing brain.

### Schizophrenia risk genes disrupt forebrain gene expression network

Using 189 dorsal and ventral forebrain organoids, we performed 90 bulk RNA-seq with 3-5 samples per KO group from 2-3 batches and each sample used 2-3 organoids and each batch includes 2-3 control samples at 6 weeks **(Figure 2a, b)**. The average percentages of co-upregulated and co-downregulated genes were 11.64% (95% confidence interval, 8.21%-15.06%) and 16.55% (95% confidence interval, 11.93%-21.16%) in both DFOs and VFOs, which indicated that these 11 schizophrenia risk genes play different roles in excitatory neurons and inhibitory neurons **(Figure S5a, b** and **Table S2)**. Rank-Rank Hypergeometric Overlap (RRHO) analysis further confirmed that the dysregulation of expression profiles in DFOs and VFOs with the same KO risk gene are different **(Figure S5c, d)**. Furthermore, based on the results of differential expression analysis from bulk RNA-seq, we conducted RRHO analysis between any two risk genes. Strikingly, gene expression profiles of DFOs, with different mutations including *CRHR1^−/−^*, *GRM1^−/−^*, *MAPT^−/−^*, *PDE4B^−/−^*, *PCCB^+/−^*, *SF3B1^+/−^* and *THOC7^+/−^*, showed a significant overlap between each other, however this similarity was not observed in VFOs **(Figure 2c)**. In VFOs, only *CLCN3^−/−^* presented significantly overlaps with *GPM6A^−/−^*and *THOC7^+/−^* **(Figure 2c)**. WGCNA module analysis demonstrated that 7 (*i.e.*, darkgrey, floralwhite, lightcyan, lightsteelblue1, plum2, salmon, skyblue) out of 22 gene expression modules in DFOs are significantly related to more than three tested risk genes, but only 1 (*i.e.*, darkgrey) out of 19 expression modules in VFOs is regulated by 3 risk genes, further indicating that these risk genes have more similar effects on DFOs, but not in VFOs **(Figure S5e, f)**.

**Figure 2.**
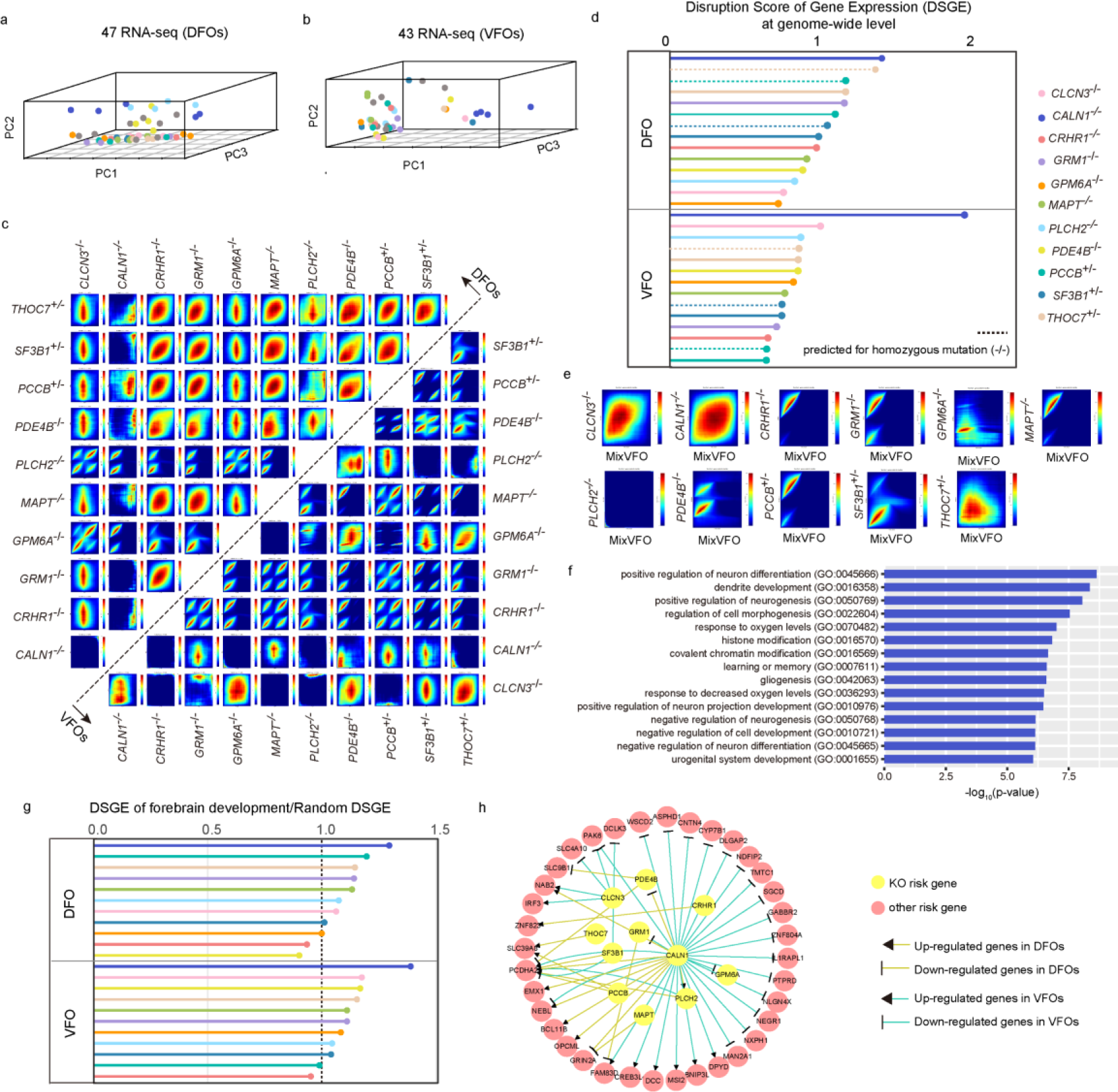
Schizophrenia risk genes regulate forebrain gene expression network. **a, b**, Top three PCs of gene expression of DFOs (**a**) and VFOs (**b**) at 6 weeks. Each group included 3-5 samples from 2-3 batches and each sample used 2-3 organoids for RNA-seq. PCs, principal components. **c**, The Rank-Rank Hypergeometric Overlap (RRHO) maps between any two genes in DFOs or VFOs. Signals in top right quadrant and bottom left quadrant indicate an overlap of down-regulated and up-regulated genes found in both KO genes, respectively. **d,** The disruption score of gene expression (DSGE) for each mutation based on z-score of differentially expressed analysis of 6-week organoids at genome-wide level, of which, including the predicted DSGE of homozygous mutation for three heterozygous mutations (*i.e.*, *PCCB^+/−^*, *SF3B1^+/−^* and *THOC7^+/−^*). **e,** The RRHO maps between the MixVFOs and VFO with single gene knockout. **f,** Top 15 GO terms related with biological process enriched by differentially expressed genes in MixVFOs. **g**, The ratio between DSGD of forebrain development and random DSGE in every KO organoid. This statistic value with greater than 1 indicates the corresponding gene affects forebrain development. **h**, 40/106 known schizophrenia risk genes are dysregulated in KO organoids.

To quantify the knock-out effect of each single gene, we calculated the disruption score of gene expression (DSGE) for every mutation based on z-score of differentially expression analysis of 6-week-old organoids at genome-wide level, of which, including the predicted DSGE of homozygous mutation for three heterozygous mutations (*i.e.*, *PCCB^+/−^*, *SF3B1^+/−^* and *THOC7^+/−^*) (**Figure 2d**). The results showed that *CALN1* lead to the strongest degree of dysregulated gene expression profile at the genome-wide level, especially in VFO (**Figure 2d**). Besides, we combined all validated clones in a pooled village-in-a-dish with equal proportions of knockout cells per gene and then induced mix VFOs (MixVFOs). At week 6, using bulk RNA-seq and RRHO analysis combination of single gene knockout, we found that the dysregulation of gene expression in MixVFOs was mostly mediated by *CALN1*, followed by *CLCN3* and *THOC7*, which further confirmed *CALN1^−/−^*inducing the largest effects of all 11 risk genes (**Figure 2e**). Gene Ontology (GO) enrichment analysis established differentially expressed genes in MixVFOs enrich in pathways associated with neuronal differentiation, histone modification, covalent chromatin modification, gliogenesis and so on (**Figure 2f**).

For further evaluating the impacts of these risk genes on forebrain development, we collected 379 genes associated with forebrain development and calculated DSGE of these genes in different knock-out DFOs and VFOs. Next, we assessed the DSGE of random 379 genes across genome in 200 random samples and the average value was used to compare with DSGE of forebrain development. The results display that the ratio of DSGD of forebrain development and random DSGE in most of knock-out DFOs and VFOs are greater than 1, suggesting these risk genes significantly affect forebrain development (**Figure 2g**). And then we furtherly illustrate the specific dysregulated genes associated with forebrain development in each knock-out DFOs or VFOs **(Figure S5g-j)**. Notably, regardless of dorsal or ventral, and up- or down-regulated, knockout of *CALN1* can lead to the massive gene dysregulation of forebrain development. In addition to genes associated with forebrain development, we found that 40 out of 106 schizophrenia risk protein-coding genes were involved in abnormal expression in these 11 gene KO organoids, of which including protocadherin alpha 2 (*PCDHA2*) dysregulated in 5/11 KO genes and its roles in neuronal development involved in psychiatric disorders were proven by several studies ^12,28,29^ **(Figure 2h)**. Notably, 34 of 106 schizophrenia risk genes are significantly dysregulated in *CALN1^−/−^*organoids. (**Figure 2h**).

Enrichment analysis showed that the up-regulated genes in DFOs with *CALN1^−/−^* were mainly enriched in forebrain pattern specification, neuron projection and forebrain regionalization, while down-regulated genes were involved in extracellular matrix organization and plasma membrane **(Figure 3a** and **Figure S6a)**. Correspondingly, the up-regulated genes in VFOs with *CALN1^−/−^* have effects on cell cycle and pattern specification process, but down-regulated genes were enriched in synapse organization and assembly and synaptic transmission (**Figure 3b** and **Figure S6b)**. Bulk RNA-seq and immunostaining analysis revealed that the expression of *PAX6* in DFOs and *NKX2-1* in VFOs with *CALN1^−/−^* are significantly higher than wild type **(Figure 1f, g** and **Figure 3c, d)**. At the same time, we found that there is a significant reduction of *DCX^+^* young neurons in DFOs **(Figure S6c)**. Besides, the expression of glial cell marker *S100B* is significantly lower in both *CALN1^−/−^* DFOs and VFOs **(Figure 3e, f)**. These data further verified that *CALN1* regulates the differentiation of the dorsal and ventral neural progenitor cells. Taken together, the collected results indicated that knockout of 11 schizophrenia risk genes lead to different degrees of effects in both human DFOs and VFOs, and have more similar effects on DFOs compared with VFOs. Among them, *CALN1* contributes the most comprehensive and severe disturbance of human forebrain development.

**Figure 3.**
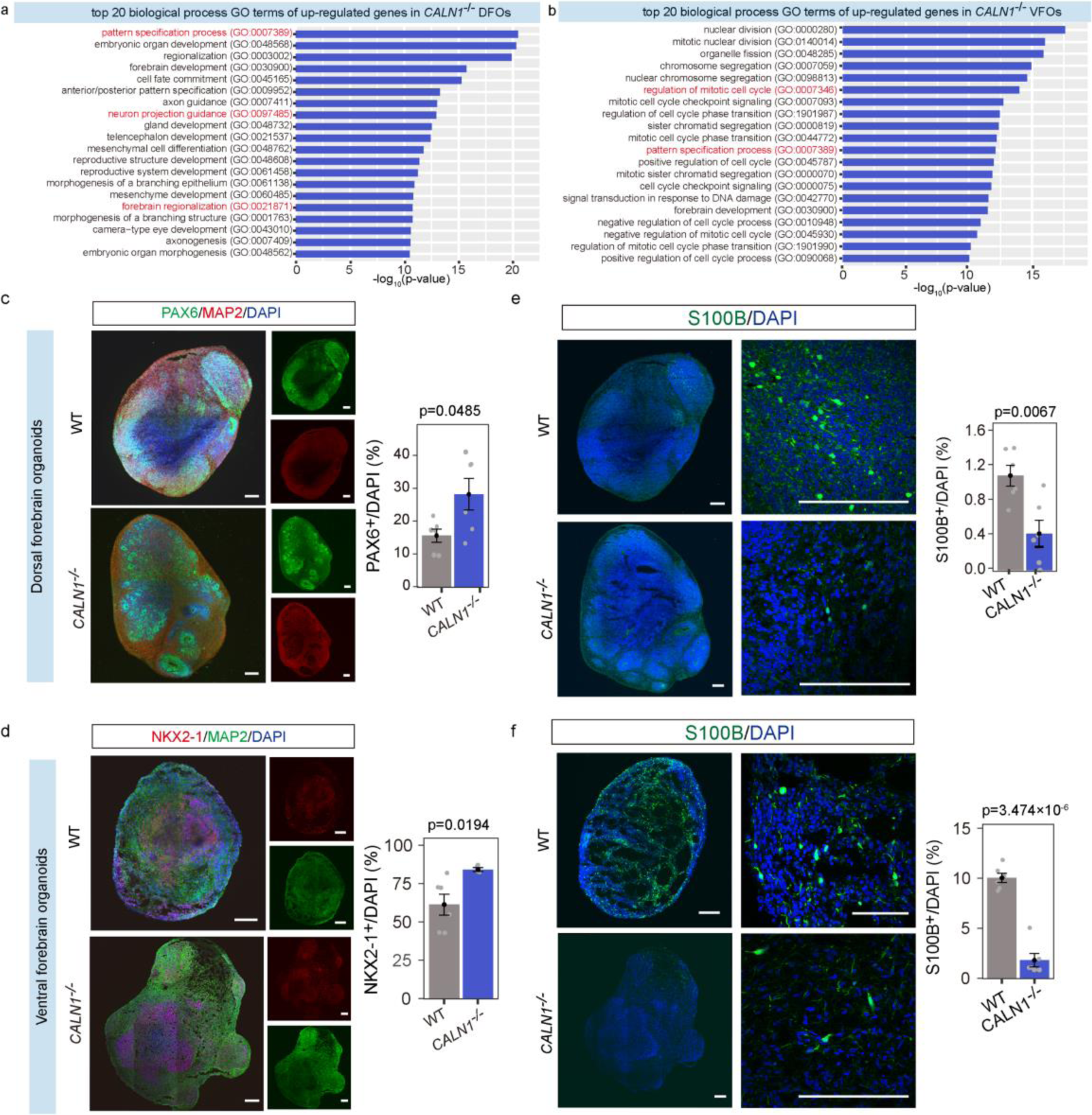
Knockout of *CALN1* impacts the early development of forebrain organoids at 6 weeks. **a, b,** Top 20 GO terms related with biological process enriched by up-regulated genes in *CALN1^−/−^* DFOs (**a**) and VFOs (**b**). **c-f,** Immunostaining of specific cell markers, including PAX6 (**c**) and S100B (**e**) in *CALN1^−/−^* DFOs and NKX2-1 (**d**) and S100B (**f**) in *CALN1^−/−^* VFOs. The p-values were calculated using two tailed t-test. scale bar, 200 μm.

### Schizophrenia risk genes affect specific cell population and development trajectories

We profiled 56 DFOs and VFOs with wild type and 11 KO genes at 12 weeks using single-cell RNA-sequencing and 4-6 organoids were used per KO gene. A total of 249,430 qualitied cells (44,899 cells with wild type, and 204,531 cells with 11 KO genes) were obtained for further analysis **(Figure 4a, c)**. We identified 14 and 13 cell populations according to known marker genes for DFOs and VFOs with wild type, respectively **(Figure S7a-d)**. The diffusion maps showed that both DFOs and VFOs have two distinct groups of cells (*i.e.*, neurons and glial cells) that differentiated from radial glial cells **(Figure S7e, g)**. The branching dendrogram structures, which were built based on cell pseudotimes, further depicted timing characteristics of each neuron and glial cell population **(Figure S7f, h)**.

**Figure 4.**
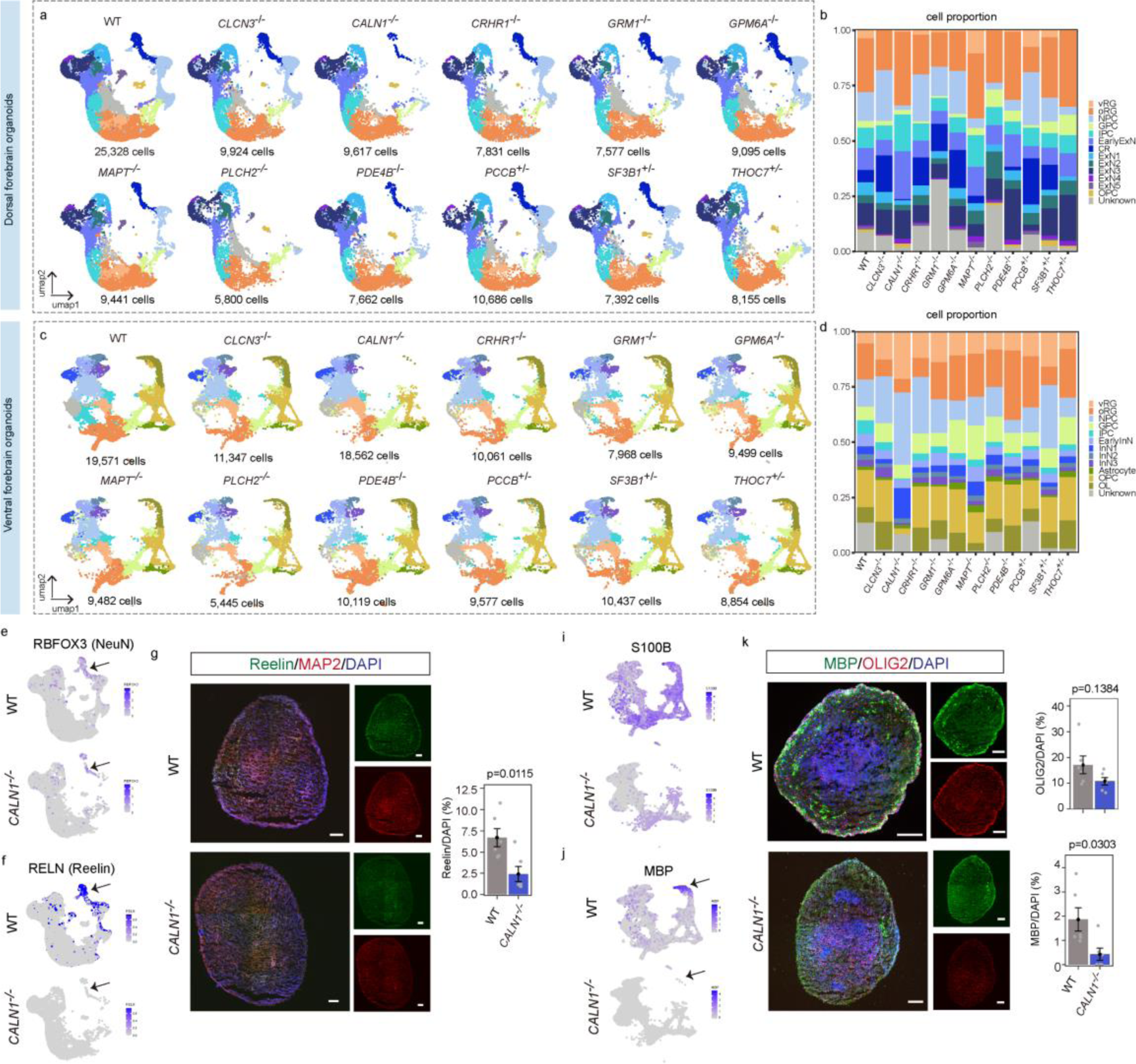
Schizophrenia risk genes affect specific cell population in forebrain organoids at 12 weeks. **a, c**, The UMAP plots of scRNA-seq for each KO DFO (**a**) and VFO (**c**) at 12 weeks. UMAP, Uniform Manifold Approximation and Projection. **b, d**, The proportion of each cell population in scRNA-seq for each KO DFO (**b**) and VFO (**d**). **e, f**, The reduction of mature neurons with expression of *RBFOX3* and *RELN* in scRNA-seq of *CALN1^−/−^* DFOs. **g,** Immunostaining of Reelin in *CALN1^−/−^*DFOs. **i, j**, The reduction of glial cells with expression of *S100B* and *MBP* in scRNA-seq of *CALN1^−/−^* VFOs. **k,** Immunostaining of OLIG2 and MBP in *CALN1^−/−^* VFOs.

Next, we used scRNA-seq data of DFOs/VFOs with wild type as reference to identify the cell type of each qualified KO organoid and counted the proportion of various cell populations in all organoids. In DFOs, the EarlyExN cells, which identified based on *DCX^+^* and *PAX6^−^*, with *CALN1^−/−^* have robustly increased 50% compared with wild type **(Figure 4a, b** and **Table S3)**. Immunofluorescence staining further confirmed that the expression of DCX is significantly higher in 12-week-old DFOs with *CALN1^−/−^*(**Figure S6c**). At the same time, based on scRNA-seq and immunofluorescence staining, we found that there is a reduction of mature neurons with expression of NeuN in *CALN1^−/−^* DFOs **(Figure 4e** and **Figure S6d)**, and further analysis indicated that the missing mature neurons are may predominantly *RELN^+^* cells (**Figure 4f**). And then immunofluorescence staining also confirmed the reduction of Reelin at the protein level in *CALN1^−/−^* DFOs (**Figure 4g**). Besides, the percentage decrease of Cajal-Retzius (CR) cells in DFOs with *PLCH2^−/−^*, *PDE4B^−/−^* and *THOC7^+/−^* at 12 weeks is also more than 50% compared with wild type **(Figure 4a, b** and **Table S3)**, which suggesting these risk genes may play important roles in the development of mature neurons in the superficial layer 1 (L1) of the cortex. The increased percentages of neural progenitor cells (NPC) in VFOs with *CLCN3^−/−^*, *CALN1^−/−^*, *CRHR1^−/−^* and *SF3B1^+/−^* are up to 50% compared with wild type **(Figure 4c, d** and **Table S3)**. InN1 cells of *SST^+^* in VFOs with *CALN1^−/−^*significantly increase, which was supported by both scRNA-seq and immunostaining data **(Figure 4c, d, Figure S6e** and **Table S3)**. Notably, the expression of glial cell marker is significantly reduced in *CALN1^−/−^* VFO (**Figure 4i**), and both scRNA-seq and immunostaining furtherly confirmed that the development of oligodendrocytes (OL) (*i.e.*, *OLIG2^+^* & *MBP^+^*) is severely inhibited (**Figure 4j-k**).

We also estimated cell pseudotime using radial glial cells as original cell population and built developmental trajectories of each organoid. Compared with wild type, we found that the pseudotime densities of different cell lineages in different KO organoids are distinctive **(Figure S8a, b)**. To quantify the maturation of cell population, one-tailed Kolmogorov-Smirnov tests were performed on the start and end point of trajectory, respectively. Maturational delay of CR cells were observed in DFOs with *CALN1^−/−^* and *PDE4B^−/−^*, while accelerated maturation of CR population with *CLCN3^−/−^*, *CRHR1^−/−^* and *PCCB^+/−^* (p-value<5×10^−15^, **Figure S8c**). Meanwhile, ExN2, ExN3 and ExN4 neuron populations are more mature in DFOs with *CALN1^−/−^* and *PDE4B^−/−^*(**Figure S8c**). Strikingly, the whole neuron lineages (*i.e.*, EarlyInN, InN1, InN2 and InN3) in VFOs with KO genes showed increased distributions towards the end point of the trajectory, indicating the accelerated maturation of neurons **(Figure S8d)**. The OPC population (*i.e.*, *PDGFRA^+^* and *MBP^−^*) is significantly immature in VFOs with *CALN1^−/−^* **(Figure S8d)**. To examine the mechanisms for abnormal development of neurons, we performed differential expression analysis for each neuron lineage. Differential expression analysis of neuron-related clusters in DFOs and VFOs showed that a total of 70 and 73 genes are abnormally expressed, respectively, and each KO organoids displayed different expression patterns in various neuron clusters **(Figure S8e, g)**. GO enrichment analysis found that these differentially expressed genes (DEGs) are key genes related to forebrain development, neurogenesis and so on **(Figure S8f, h** and **Table S4)**. Taken together, these results indicated that knockout of these schizophrenia risk genes have more similar effects on superficial layer neurons in DFOs, while the roles in VFOs are more diverse and specific. Pseudotime and differentially expressed analysis showed that neuronal dysregulation is mediated by different genes in these knock-out organoids and *CALN1^−/−^* leads to the delayed neuronal maturation and the deficit of gliogenesis.

### Schizophrenia risk genes affect spontaneous neural circuit activity

The abnormalities in gene expression network, specific cell population and developmental trajectories in these KO organoids prompted us to investigate their neural functional activity. We analyzed spontaneous neuronal activity of 145 DFOs and VFOs at 12 weeks including wild type and KO genes by multielectrode array (MEA). There were 61,654 and 88,850 spikes generated by 962 and 893 active electrodes from DFOs and VFOs, respectively **(Figure 5a, b** and **Figure S9a)**. We found that the alternations of spontaneous neuronal activity are different in DFOs and VFOs. In addition to *MAPT* and *PDE4B*, significantly longer inter-spikes intervals (ISIs) were observed in other KO DFOs, while shorter ISIs in multiple KO VFOs apart from *CALN1*, *PLCH2* and *SF3B1* **(Figure 5c-f)**. At the level of active electrodes, 7/11 KO DFOs (*i.e.*, *CALN1^−/−^*, *GRM1^−/−^*, *GPM6A^−/−^*, *PLCH2^−/−^*, *PDE4B^−/−^*, *PCCB^+/−^* and *THOC7^+/−^*) induce significantly lower firing rates, which indicate weaker neuron activity (**Figure 5g**). Notably, knockout of *GRM1*, results in weaker neuronal activity and neuron-neuron connections in DFOs, while abnormally higher synchronized firings were detected in VFOs (**Figure 5g-j** and **Figure S9a**). Besides, DFOs with *MAPT^−/−^* have stronger individual neuron activity but weaker neuron-neuron connections (**Figure 5g, h**). The significantly longer ISIs, lower firing rates and weaker synchronization were observed in both DFOs and VFOs with *CALN1^−/−^* at 12 weeks, indicating both neuronal activity and connections are affected by *CALN1* (**Figure 5c-j**). Taken together, these results indicated that most of these risk genes may play roles in the inhibition of excitatory neuronal activity in DFOs and the promotion of inhibitory neurons in VFOs. Among them, *CALN1* exhibits strong inhibitory effects on both neurons.

**Figure 5.**
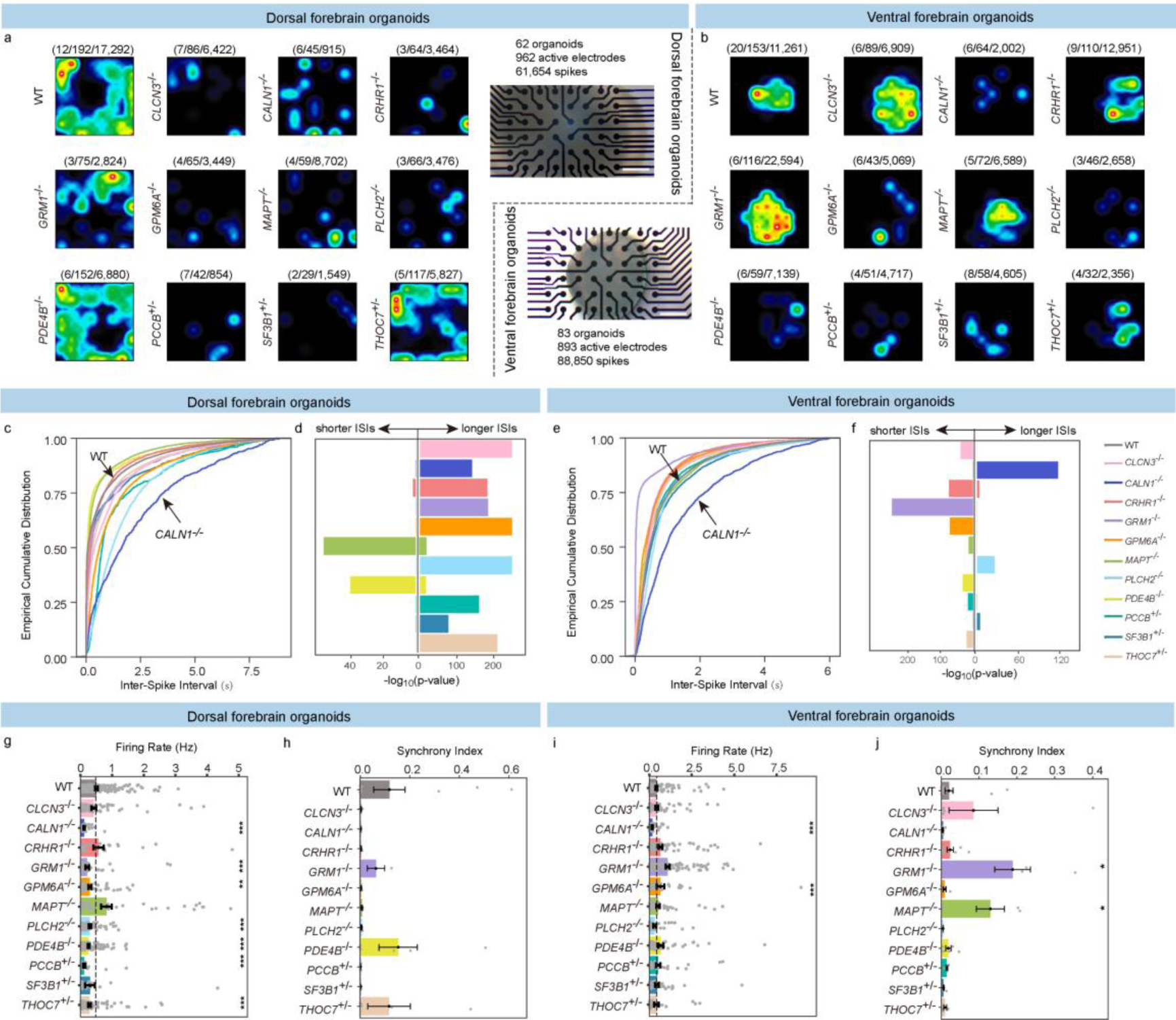
The multielectrode array (MEA) recording and analysis of knock-out dorsal and ventral organoids. **a, b**, Example of electrode-activity heatmaps in DFOs (**a**) and VFOs (**b**) with knock-out gene and wild type. The number of organoids, active electrodes and spikes were labeled in turn above each heatmap. Scale bar, 500 μm. **c, e**, Empirical cumulative distribution plots of Inter-Spike Intervals (ISIs) for DFOs (**c**) and VFOs (**e**). **d, f**, Knock-out DFOs (**d**) and VFOs (**f**) have significantly longer ISIs (right) or shorter ISIs (left) compared with wild type (one tail Kolmogorov-Smirnov test). **g, i**, Firing rates of each electrode in DFOs (**g**) and VFOs (**i**). **h, j**, Synchrony indexes of each DFO (**h**) and VFO (**j**). The two-tailed t-test was used in **g-j**. Data are presented as mean ± SE. ***, p-value < 0.001. **, p-value < 0.01. *, p-value < 0.05.

### Common pathways are disturbed in both *CALN1^−/−^* forebrain organoids and cerebral cortex of *Caln1^−/−^* E17.5 mice

Given the remarkable dysregulation of key developing genes and neuronal functional activity in *CALN1* KO forebrain organoids, we next constructed *Caln1^−/−^* transgenic mice (**Figure S10a-c**). To furtherly explore the effects of *Caln1* in the developing brain, we performed bulk RNA-seq using cerebral cortex tissues from E17.5 *Caln1^−/−^* mice. Through differentially expressed analysis and GO enrichment analysis, pathways associated with neuron to neuron synapse, neuron migration, glial cell differentiation, axonogenesis and so on are significantly dysregulated in both *CALN1^−/−^* organoids and *Caln1^−/−^* mice (**Figure 6a**). Besides, 67 DEGs identified in DFOs and 146 DEGs identified in VFOs were also detected in *Caln1^−/−^*mice (**Figure 6b, c** and **Table S2**). Notably, *LIN28A/Lin28a* and *PEG3/Peg3* are significantly up-regulated in knock-out organoids and mice (**Figure 6d**), which may be involved in the maintenance of neural progenitor cells and disturbing neural cell differentiation^30–32^.

**Figure 6.**
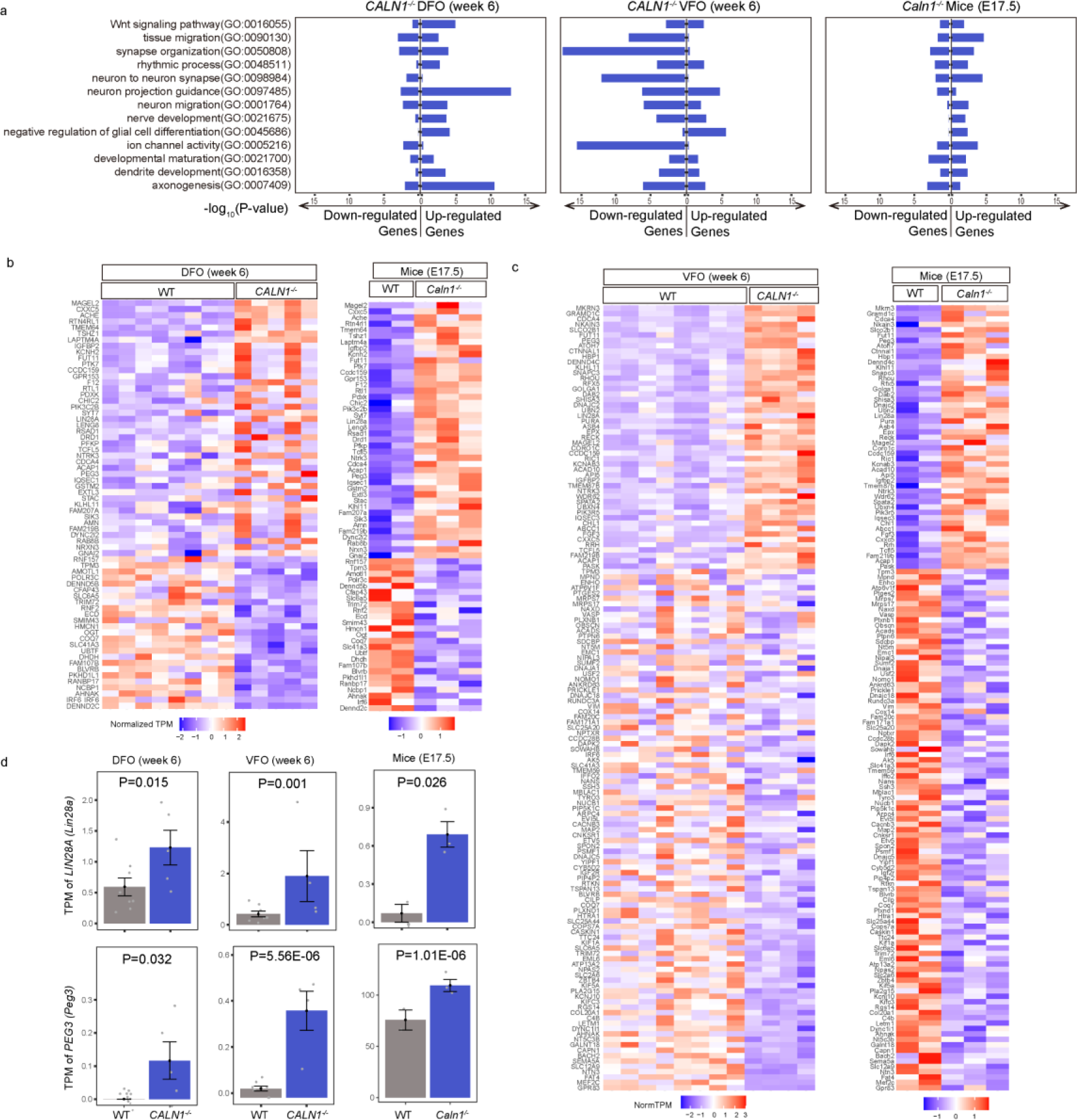
Common pathways disturbed in both *CALN1^−/−^* forebrain organoids and cerebral cortex of *Caln1^−/−^* E17.5 mice. **a,** The common GO terms in both *CALN1^−/−^* organoids and *Caln1^−/−^* mice. **b,** Common differentially expressed genes (DEGs) in both *CALN1^−/−^* DFOs and *Caln1^−/−^*mice. **c,** Common DEGs in both *CALN1^−/−^* VFOs and *Caln1^−/−^* mice. **d,** *LIN28A/Lin28a* and *PEG3/Peg3* are significantly up-regulated in both *CALN1^−/−^* forebrain organoids and *Caln1^−/−^* mice.

### *Caln1^−/−^* cortical neurons exhibit spontaneous abrupt depolarization burst firing

Except effects of early development, we also investigated electrophysiological properties of matured neocortical neurons in *Caln1^−/−^* mice (**Figure 7a-f**). We performed whole-cell current-clamp recording from layer-V pyramidal cells (PCs) in acute slices of the prefrontal cortex of adult mice (**Figure 7a**). Surprisingly, in *Caln1^−/−^* mice, 31.6% recorded cells (12/38, N = 3 mice) showed abrupt burst spiking (ABS) from their resting membrane potential (*V*_m_) (**Figure 7b-d**), a phenomenon rarely observed in prefrontal cortical neurons of WT mice (n = 30, N = 3 mice). The ABS occurred randomly during the recording with duration ranging from 0.04 s to 725 s (**Figure 7f**). Close examination of the abrupt burst revealed a train of action potentials (with a frequency of 4.1 ± 0.71 Hz, n = 40 abrupt bursts) riding on top of a depolarization plateau (**Figure 7f**). To examine whether ABS neurons possess distinct intrinsic membrane properties, we compared the electrophysiological properties of ABS and non-ABS neurons of *Caln1^−/−^* mice with those of WT mouse neurons. We found no significant difference in their resting *V*_m_ (WT, –66.4 ± 0.6; ABS, –65.5 ± 1.35; non-ABS, –66.7 ± 0.9 mV; P > 0.05 for all comparisons, Mann-Whitney test), input resistance (WT, 128.2 ± 10.2; ABS, 138.7 ± 24.0; non-ABS, 139.6 ± 15.4 mV, P > 0.05 for all comparisons, Mann-Whitney test;), and membrane time constant (WT, 21.1 ± 1.6; ABS, 20.6± 3.3; non-ABS, 20.1 ± 2.1 ms, P > 0.05 for all comparisons, Mann-Whitney test). A slight increase in the neuronal excitability was observed in ABS neurons as compared to those of non-ABS neurons and WT neurons, as reflected by the upward shift of the input-output (*I-F*) curve of ABS cells (**Figure 7e**). In consistent, we also found a slight decrease in the threshold current (rheobase) for action potential generation (WT, 105.7 ± 6.9; ABS, 96.4 ± 11.9; non-ABS, 108.8 ± 7.8 pA, P > 0.05 for all comparisons, Mann-Whitney test). Taken together, these results reveal that a subpopulation of *Caln1^−/−^* cortical neurons exhibits aberrant features of spontaneous random and abrupt-burst firing.

**Figure 7.**
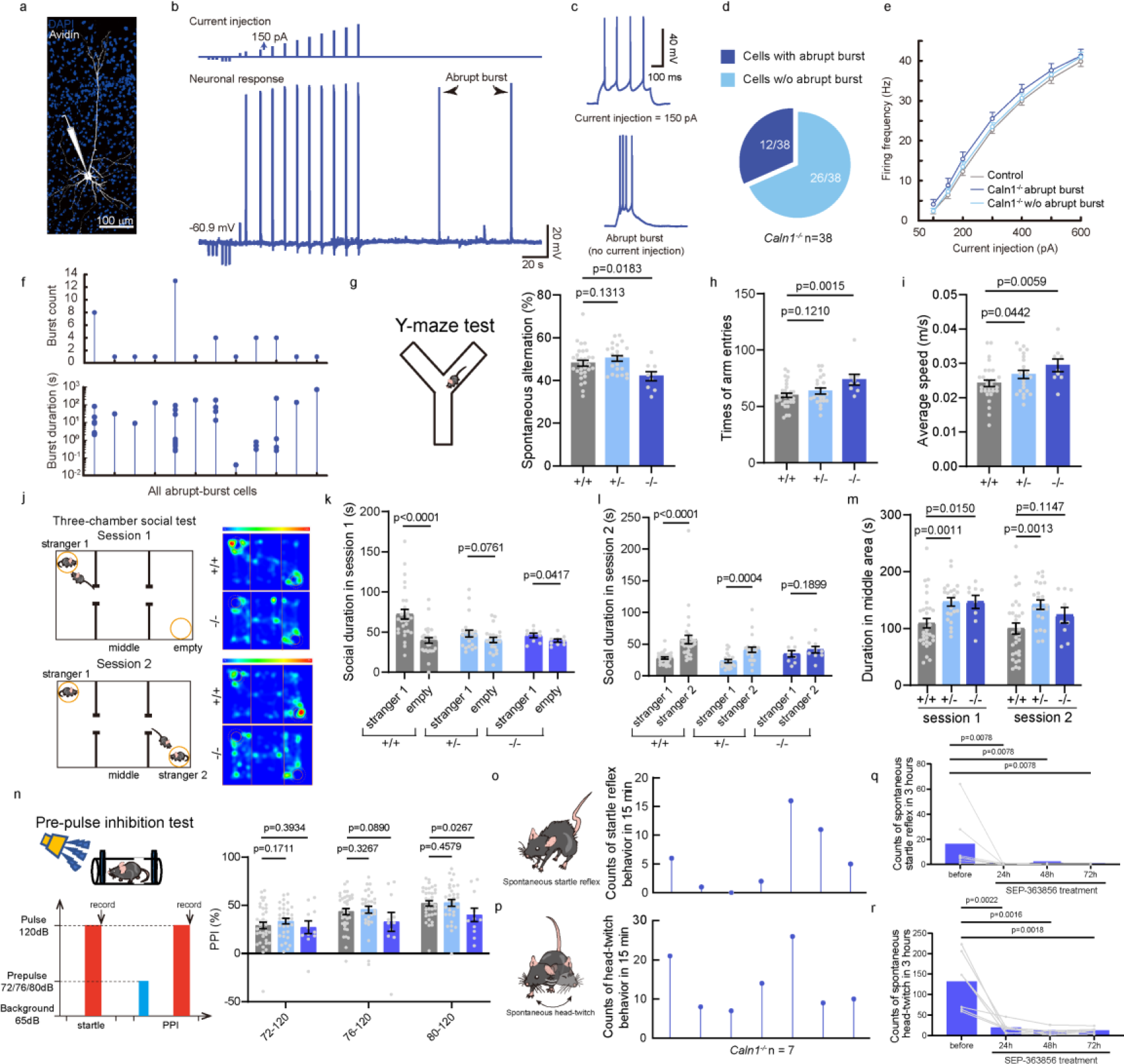
The neocortical neurons of *Caln1^−/−^* mice displayed abrupt depolarization burst firing and *Caln1^−/−^* mice displayed schizophrenia behavior. **a,** Projection of confocal z-stack images of a layer-V pyramidal neuron that has been filled with biocytin during whole-cell recording in a prefrontal cortical slice. **b**, An example of continuous recording from a *Caln1^−/−^* cortical neuron showing membrane potential responses (bottom) to intracellular injections of step currents (top) and the occurrence of abrupt burst spiking (ABS) events from the resting membrane potential. **c**, Representative traces showing spiking responses to a current step of 150 pA (top) and the two ABS events (bottom). **d**, Percentage of ABS (dark blue) and non-ABS neurons (light blue) from *Caln1^−/−^* mice. **e**, Comparison of *I-F* curves of WT neurons, *Caln1^−/−^* ABS and non-ABS neurons. **f**, Counts of abrupt burst events in individual ABS neurons (top) and their duration (bottom). **g-i**, Y-maze test of three genotypes of mice. Comparing with wild type, *Caln1^−/−^* mice have significantly lower the percentage of spontaneous alternation (**g**), more total number of arm entries (**h**), and average speed (**i**). **j-m**, The three-chamber social test. The heatmaps of social time between WT and *Caln1^−/−^* mice (**j**), social duration with stranger mouse and empty cage in session 1 (**k**), social duration with stranger 1 and stranger 2 in session 2 (**l**) and duration in the middle area (**m**) were showed. **n**, The 120dB startle stimulus and three different pre-pulse stimuluses (*i.e.*, 72, 76 and 80 dB) were conducted in the pre-pulse inhibition (PPI) test. Data are presented as mean ± SE. The one-tailed t-test was used to calculate p-value. **o, p**, The counts of spontaneous startle behavior (**o**) and head-twitch response (**p**) within a random 15 minutes from 7 *Caln1^−/−^* mice. **q, r,** The counts of spontaneous startle behavior (**q**) and head-twitch response (**r**) before and after treatment with SEP-363856 for 3 hours from 7 *Caln1^−/−^* mice. The paired nonparametric test was used to calculate p value.

### *Caln1^−/−^* mice display schizophrenia behavior

Abnormal neuronal activities suggest that *Caln1^−/−^* mice may have behaviors deficits, and then, we conducted classical behavior tests for examining learning and memory, social ability and depression, using WT, *Caln1^+/−^*and *Caln1^−/−^* mice at 8-11 weeks. In the Y-maze test, comparing with wild type, *Caln1^−/−^* mice have significantly lower percentage of spontaneous alternation, displaying significant deficiency of short-term spatial working memory and cognition compared with wild type mice **(Figure 7g)**. Meanwhile, *Caln1^−/−^* mice showed a remarkable increase of the total number of arm entries and average moving speed, which suggested that they were more active comparing with WT mice **(Figure 7h, i)**. In the three-chamber social test, *Caln1^−/−^* mice appeared weaker social preference between stranger 1 mice and empty cage than WT mice in session 1, along with spending more time in the middle area, and in session 2 *Caln1^−/−^* mice showed no significantly social tendency to stranger 2 mice compared with stranger 1 mice, which indicated an impaired sociability and a decreased social motivation in *Caln1^−/−^* mice **(Figure 7j-m)**. In addition, in the pre-pulse inhibition test (PPI), the *Caln1^−/−^* mice showed the significantly PPI deficits **(Figure 7n)**. In the open-field, elevated plus maze and tail suspension test, there were no significant difference observed between *Caln1^−/−^*and WT mice **(Figure S10d-g)**. In addition to these classic behavioral paradigms, we surprisingly observed that *Caln1^−/−^* mice from about 4 weeks old appeared spontaneous startle behavior (**Figure 7o** and **Video S1**) and spontaneous head-twitch response (**Figure 7p** and **Video S2**), which were considered as hallucination-like behavior in human^20–25^. Both specific behaviors occur up to 16 and 26 times within 15 minutes during the onset of the abnormality (N=7, **Figure 7o, p**). Moreover, the frequencies of these unusual spontaneous startle behavior and head-twitch behavior significantly reduce in 3 hours after treatment with SEP-363856, an investigational antipsychotic drug (N=7, **Figure 7q, r**)^33^. In summary, these findings verified that *Caln1^−/−^* mice appear schizophrenia behaviors including the defects of spatial memory, cognition, social ability and pre-pulse inhibition, as well as unusual spontaneous startle behavior and head-twitch response similar to hallucination in human, which could be treated by antipsychotic drug SEP-363856 (**Figure S11**).

## Discussion

Schizophrenia is considered as to be polygenic psychiatric disorder^34,35^ and previous studies have reported that some risk genes disrupt human developing neuronal and glial cells in schizophrenia, such as *PCDHA* ^12^, *C4A* ^36,37^, *NRXN1*^18^, *DISC1*^38^ and *MAD1*^39^. There are 106 protein-coding risk genes identified by fine-mapping and functional genomic analysis based on the largest GWAS of schizophrenia so far^6^. However, due to the difficulty of quantifying the contribution to disease of every genetic variant or risk gene in complex genetic structures of patients, and to understand their pathogenic mechanisms in human developing brain, we established the strategy of single-gene-knockout-precise-dorsal/ventral-forebrain-organoids (SKOPOS) including both excitatory and inhibitory neuron system (**Figure S11**). Based on knock-out excitatory dorsal and inhibitory ventral forebrain organoids, our study simultaneously and comprehensively explored the impacts of 11 schizophrenia risk genes on developing human brain from gene expression profiles, specific cell population and neural functional activity. According to bulk RNA-seq, we measured the disruption score of gene expression at both genome-wide and forebrain development levels and discovered that although these risk genes disrupt gene expression network to varying degrees, most of them have significantly higher impact on forebrain development than random events. No specific cell population showed a complete consistent increase or decrease across all 11 risk genes, but dysregulated genes in knock-out neurons were found significantly enriched in the pathway associated with forebrain development. Besides, MEA analysis suggested that inhibitory neural activity is enhanced in most of these knock-out organoids. Remarkably, among those multiple schizophrenia risk genes, *CALN1* induces the most severe perturbation of gene expression network, cell population and neural activities in human developing forebrain (**Figure S11**). The collected results indicate that different risk genes may lead to similar functional consequences through different molecular pathways and cell populations, which may explain why patients with similar clinic phenotypes respond differently to the same antipsychotics^40,41^. Considering the fact brain organoids are unable to mimic patients’ behavioral phenotypes, here, we constructed *Caln1* knock-out mice (**Figure S11**). Including the spontaneous abrupt burst spiking in cortical neurons and the defects of the defects of spatial memory, cognition, social ability and pre-pulse inhibition, *Caln1* KO mice surprisingly displayed spontaneous startle behavior and head-twitch response, which has been reported as a hallucination-like behavior in human^20–25^. And furtherly we found the frequencies of these unusual behaviors significantly reduce after treatment with SEP-363856, an investigational antipsychotic drug. It is known that hallucination is one of typical symptoms in schizophrenia patients and is experienced by 60%-80% of patients^42^, which may bring severe damage to patients, for instance, leading to suicidal ideation and suicide attempt^43^. In the previous studies, the deficit of prepulse inhibition (PPI) of acoustic startle reflex, as one of typical behaviors in schizophrenia patients, has often been used to model schizophrenia-like behavior in rodents. According to our knowledge, *Caln1^−/−^* mice is first mutant mouse model exhibiting spontaneous startle behavior and head-twitch response, which may be as a novel animal model to study the molecular mechanism of schizophrenia and hallucination, and it would be interesting to investigate how *Caln1* causes hallucination, as well as its contribution to schizophrenia.

Our study has limitations. First, each knock-out cell line used in this study only includes one gene mutation, and lacks multigenic co-perturbations like in patients. Further studies should explore the impact of multiple genes in the same cell line and whether the combination of different genes has a greater effect than a single gene. Additionally, brain organoids are good models for exploring the role of early brain development, but their impact on the late development stage, especially in adulthood, is limited, which makes it difficult to explain how a link is made between early developmental defects and behavioral abnormalities in adulthood. We think that it is necessary to construct KO animal model of key risk gene to verify the disease associated behaviors.

Polygenic risk scores (PRS) based on genomic information are often used to assess the person’s chances of suffering a complex disease^35^, including schizophrenia^44,45^. However, a person’s PRS of schizophrenia is calculated only based on the presence or absence of multiple genomic variants, without taking the strength of disruption by associated mutations or genes in the nervous system into account. Our parallel studies of multiple single gene provide important gene disruption and interaction score of functions in both excitatory and inhibitory neural system and can contribute to more accurate person’s risk assessments. Besides, we firstly discovered the roles of risk gene *CALN1* in human brain organoids and rodents, and the established *Caln1^−/−^* mouse may be a novel animal model of SCZ and can offer a promising therapeutic target as well as contribute to develop new precision drugs of schizophrenia in the future.

## Materials and Methods

### GWAS Summary Statistics of Schizophrenia Risk Genes Loci

We retrieved the summary statistics of SNPs located in 11 tested risk genes loci from the latest schizophrenia GWAS which based on 74,776 cases and 101,023 controls from European, East Asian, African American and Latino ancestry (accessed at https://www.med.unc.edu/pgc/download-results/scz/) ^6^. The most significant SNP in every region was identified as the index SNP and linage disequilibrium of other SNPs were calculated based on the genotype data of European population from the 1000 Genomes Project.

### Guide RNAs design

Guide RNAs (gRNA) were designed using the tool named Benchling (accessed at https://www.benchling.com/) and the optimal site was selected based on the principle that maximizing on-target efficiency and minimizing off-target probability in exon regions for creating a protein truncation in the early stage of protein translation. To knock out eleven schizophrenia risk genes, in this study, selected gRNAs were as follows, 5’-TCTGAGCAGCTGTTCCATAG-3’ (AGG) (for *CLCN3*), 5’- ACAGCTGGCTAATATCTCCG-3’ (TGG) (for *CALN1*), 5’- GTACACCGACTACATCTACC-3’ (AGG) (for *CRHR1*), 5’- CACGTTGGATAAGATCAACG-3’ (CGG) (for *GRM1*), 5’- GGAAGGTTTCTTCACAACTG-3’ (CGG) (for *GPM6A*), 5’- CTGGTGCTTCAGGTTCTCAG-3’ (TGG) (for *MAPT*), 5’- TGACGAGGCCGACAAGAACG-3’ (GGG) (for *PLCH2*), 5’- TCTCAGAGATGAGCCGATCA-3’ (GGG) (for *PDE4B*), 5’- ATTGGGCTGAATGACTCTGG-3’ (GGG) (for *PCCB*), 5’- TGTCCACTCGAACACACAGA-3’ (CGG) (for *SF3B1*), 5’- ACAGCGTGCTCAGCATACGT-3’ (TGG) (for *THOC7*). Besides, we designed a gRNA (*i.e.*, 5’-GGTCTTCCCGACGATGACGC-3’) that doesn’t target human genome as positive control.

### hESCs Culture

The WilCell Research Institute’s H9 (WA09) hESCs were maintained in NutriStem^®^ hPSC XF Culture Medium from Biological Industries (cat. #05-100-1A) on 35mm dishes coated with 1% Matrigel (Corning, cat. #354230). hESCs were passaged using Gibco^TM^ TrypLE^TM^ Express Enzyme (cat. #12604021) and proved mycoplasma negative. Below passage 45 of hESCs were used to knock out schizophrenia risk genes and then less 10 passages of knock-out hESCs were induced to be organoids. All hESCs were cultured in a 5% CO_2_ incubator at body temperature 37 ℃.

### CRISPR-editing human embryonic stem cell lines

Each gRNA was cloned into the LentiCRISPRv2 vectors including Cas9 sequence (Addgene #52961), and then were used to lentivirus package based on calcium phosphate method along with plasmids psPAX2 (Addgene #12260) and pMD2.G (Addgene #12259). The 200 μl lentivirus supernatant was added to 50%-60% confluent hESCs and hESCs were selected by adding three-day 0.5μg/ml puromycin after 48 h of infection. We picked more than 5 puromycin-resistant positive cell colonies per gene for proliferation, and Sanger sequence was used to confirm the genotype each colony. Immunofluorescence staining of OCT4 and SSEA4 were performed to test whether the pluripotency of hESCs with knock-out risk gene altered.

### Generation of dorsal and ventral forebrain organoids

To generate DFOs and VFOs, we modified the previous methods in ref.^12,46^. Specifically, on day 0, hESCs (as negative control), hESCs with non-human gRNA (as positive control) and hESCs with knock-out risk gene were dissociated to single cell using Gibco^TM^ TrypLE^TM^ Express Enzyme (cat. 12604021) and 5,000 cells were reaggregated in U-bottomed 96-well plates with ultra-low-cell adhesion (Corning, cat. 7007) in dorsal or ventral forebrain differentiation medium (named dDM/vDM I). Basal medium of dDM/vDM I is DMEM containing 20% Knockout Serum Replacement (KSR) (Gibco), 1% GlutaMAX (Gibco, cat. #35050061), 10 μM β-mercaptoethanol, 1% penicillin-streptomycin (Gibco, cat. #15140122). For the first week, 10 μM TGFβ inhibitor SB431542 and 0.1 μM BMP inhibitor LDN193189 were added to the dDM/vDM I and additional 0.1 μM SAG and 5 μM IWP2 were added to vDM I. Besides, from day 0 to day 2, 10 μM ROCK inhibitor Y-27632 was added to the dDM/vDM I and cell was incubated in ultra-low 96-well plates in a static mode. After day 3, the aggregates were transfer to six-well plates (Corning) and cultured on the orbital shakers at 70 r.p.m. At the end of the first week, we embedded every organoid with 5 μl Matrigel (Corning, cat. #354230).

The medium used in dorsal forebrain organoids from week 2 to 3 was named dDM II and after 3 weeks was named dDM III. And the mediums used in ventral forebrain organoids include vDM II (for week 2), vDM III (for week 3) and vDM IV (after 3 weeks). vDM II was no longer added SB431542 and IWP2 compared to vDM I. Basal medium of dDM II, vDM III and vDM IV is DMEM/F12 (Gibco) supplemented with 2% B27 (Gibco). dDM II contained 20 ng/μl bFGF (PeproTech, cat. #AF-100-18B) and 20 ng/μl EGF (R&D Systems, Cat. #236-EG) and vDM III contained 0.1 μM SAG and 50 ng/μl FGF8 (PeproTech, AF-100-25). Content of dDM III and vDM IV were the same, that is, basal medium was supplemented with 10 ng/μl GDNF (PeproTech cat. # AF-450-10), 10 ng/μl BDNF (PeproTech cat. # AF-450-02) and 200 μM L-Ascorbic acid (Sigma-A8960).

### Immunostaining

Samples were fixed in 4% paraformaldehyde (PFA) and then cryoprotected in 30% sucrose solution overnight until sample sinks. The optimal cutting temperature medium (OCT)-embedded samples were sectioned at 25 μm thickness. Before immunostaining, cryosections were washed three times in phosphate buffered saline (PBS) and for 10 min each time. Cryosections were blocked for 30 min at room temperature in PBS with 3% Bovine Serum Albumin (BSA) and 0.3% Triton X-100 (named blocking buffer) and then incubated with primary antibodies diluted in blocking buffer overnight at 4 ℃. Primary antibodies and dilutions used in this study are listed in **Table S5**. On the next day, primary antibodies were washed three times with PBS and then cryosections were incubated with secondary antibodies and DAPI staining (1:500) for one hour at room temperature. The confocal fluorescence microscope (Olympus FV3000) was used to capture fluorescence images and images were analyzed and processed with the software ImageJ (v1.53t).

### Bulk RNA-sequencing

Total RNA was extracted using RNAiso Plus (TaKaRa, cat. #9109) according to the product manual. Quality control of RNA, the construction of cDNA library and sequence were conducted by Shanghai OE Biotech Co., Ltd. Company. The raw 150 bp paired-reads sequenced from the Illumina NovaSeq 6000 were preprocessed using Trimmomatic-0.36 and clean reads were mapped to reference genome GRCh38/GRCm38 using HISAT2 (v2.2.1)^47^. Based on gene annotation resource GRCh38.104/GRCm39.109, the reads count of every gene was calculated using featureCounts (v2.0.1)^48^ and then counts were further normalized using transcripts per million (TPM). The Wald significance tests of R package DESeq2 (v1.40.2)^49^ was used for differential expression analysis. Genes with fold change (FC) > 1.5 or < 1/1.5 and p-value < 0.01 were considered as differentially expressed genes (named DEGs). Raw bulk RNA-seq data have been deposited within the Genome Sequence Archive for Human (https://ngdc.cncb.ac.cn/gsa-human/) with the accession identifier HRA005215.

### Real-time quantitative polymerase chain reaction (RT-qPCR)

To verify some gene knockout in this study, primers used in qPCR include 5’- TCTGAGCAGCTGTTCCAT-3’ (R1) and 5’-TCCAACTGCAGTGTCACCATC-3’ (R2) for *CLCN3*, 5’-GTACACCGACTACATCTACC-3’ (R1) and 5’- CCCGGGATTGACGAAGAACA-3’ (R2) for *CRHR1*, 5’- TATGGCATCCAGAGGGTGGA-3’ (R1) and 5’-CACCGGTTGATCCCATCCTT-3’ (R2) for *GRM1*, 5’-CAGGGAACCAGGTGTCTGAA-3’ (R1) and 5’- ACTCCAAAGCGTGAGATGCT-3’ (R2) for *PDE4B*, 5’- TTCCGTCTGTGTGTTCGAGT-3’ (R1) and 5’-ATCTGCTGTCACTTCCACCA-3’ (R2) for *SF3B1* and 5’-CCAACGTATGCTGAGCACG-3’ (R1)/ 5’- ATCACTTTTGCCAAAGCATCAT-3’ (R2) for *THOC7*. RNA from 6-week organoids were extracted and collected, and complementary cDNA was synthesized using HiScript III All-in-one RT SuperMix Perfect for qPCR (Vnzyme). With SYBR (Cat. #A25742), quantitative PCR was performed on the ViiA 7 Real-time PCR system (Applied Biosystem) run using StepOne software and gene expression was normalized to an endogenous β-actin control gene and compared using the ΔΔCt method.

### Rank-rank hypergeometric overlap (RRHO)

The gene lists were ranked according to p values and log_2_FC from different expression analysis between wild type and knock-out group in dorsal or ventral forebrain organoids. The R package RRHO (v1.40.0)^50^ was used to estimate the hypergeometric p values for quantifying overlap of two gene expression signatures. The values of −log_10_ (p-values) were presented in a heatmap and increasing overlap was indicated from blue to red in the heat map.

### Disruption score of gene expression (DSGE)

The mean of z-score, from protein-coding genes with a baseMean greater than 0.1 based on differentially expressed analysis, was used to assess DSGE. Besides, to predict DSGE in the condition of homozygous mutation for each heterozygous mutation, we first re-evaluated the new beta values for all genes with an FDR of less than 0.1 and then estimated the read counts that these genes should have in homozygous mutations. Finally, the re-evaluated read counts were used to DEGs analysis and DSGEs assessment.

### Gene enrichment analysis

The Gene Ontology (GO) enrichment analysis of DEGs was conducted using R package clusterProfiler (v4.2.2)^51^ based on annotated database org.Hs.eg.db (v3.17.0). The hypergeometric distributions were used to test overlap degree between DEGs and known schizophrenia risk gene set.

### Transcriptional profiles of fetal brain

The normalized expression values by bulk RNA-seq from 8-37 post-conception weeks (pcw) fetal DLPFC or VLPFC were download from the BrainSpan database (http://www.brainspan.org/). A total of 17 and 16 gene expression profiles of DLPFC and VLPFC were obtained, respectively, and every sample contain 52,376 genes.

### Weighted gene co-expression network analysis (WGCNA)

Firstly, genes with less 0.1 of mean expression value in the normalized matrices of 47 dorsal forebrain organoids, 43 ventral forebrain organoids, 17 fetal DLPFC and 16 fetal VLPFC were deleted for further analysis. Next, we constructed unsigned co-expression gene modules for these four datasets using R package WGCNA (v1.72-1)^52^ based on cleaned expression matrices, respectively. Briefly, the topological overlap matrix (TOM) was calculated based on the appropriate soft-threshold power and then a dissimilarity structure was produced by R function *hclust*. The modules with less than 0.2 dissimilarity were merged using function *mergeCloseModules* and eigengenes of every merged module were estimated using function *moduleEigengenes*. The preservation of modules identified by expression profiles of dorsal or ventral forebrain organoids in fetal DLPFC or VLPFC were calculated using function *modulePreservation* and modules with z-score > 1.96 were considered been significantly preservative. The modules with p-value < 0.05 of the first eigengene between wild type and knock-out organoids were identified significantly associated with the specific knock-out gene.

### Single-cell RNA sequencing

At week 12, individual dorsal or ventral forebrain organoid was dissociated into the single-cell suspension at 37 ℃ for 30 min using enzyme stock solution (*i.e.*, 10x EBSS (Sigma-Aldrich, cat. #E7510), 30% D (+)-Glucose (Sigma-Aldrich, cat. #G7021), 1 M NaHCO_3_ (Sigma-Aldrich, cat. #S5761), 50 mM EDTA (Sigma-Aldrich, cat. #E8008), 20 unit/ml Papain (Worthington, cat. #LS03126), 10 μg/ml DNase Ⅰ (Sigma-Aldrich, cat. #11284932001) and 180 μg/ml L-cysteine hydrochloride monochloride (Sigma-Aldrich, cat. #C7880)). After filtering with a 40 μm cell strainer (Miltenyi Biotec), the cell suspensions were centrifuged at 500 g for 5 min and resuspended in Neurobasal Medium. Further processing of samples, the construction of scRNA-seq libraries and sequence were conducted by Shanghai OE Biotech Co., Ltd. Company. Briefly, the final concentration of cell suspension was adjusted to 1,000 cells/μl and cells were loaded onto a Chromium Single Cell 3’ Chip (10x Genomics) to generate single cell Gel Beads in Emulsions (GEMs). The 10x Genomics Chromium Single Cell 3’ Library and Gel Bead Kit v3 were used to construct scRNA-seq libraries and sequencing was conducted on the NovaSeq 6000 instrument (Illumina). Raw scRNA-seq data have been deposited within the Genome Sequence Archive for Human (https://ngdc.cncb.ac.cn/gsa-human/) with the accession identifier HRA005215.

### scRNA-sequencing data pre-processing

The raw scRNA-seq reads were aligned to the human genome reference GRCh38 and reads count matrices of every sample were generated with the 10x Genomics CellRanger pipeline. And then unique molecular identifiers (UMIs) counts were further pre-processed using R package Seurat (v4.1.0)^53^. Cells expressing 10-10,000 genes and less than 10% mitochondrial content were retained and gene expression values were normalized using default parameters in the function *NormalizeData*. The top 2,000 variable genes were selected based on the *vst* method for further analysis.

### scRNA-seq cluster annotation and trajectory inference

The quality-controlled data from three samples of dorsal forebrain organoids (25,328 cells) and two samples of ventral forebrain organoids (19,571 cells) with wild type, respectively, were integrated using function *FindIntegrationAnchors* and scaled using function *ScaleData*. After dimensionality reduction with principal component analysis (PCA), the nearest-neighbor graph was constructed based on the top 20 PCs, followed by cluster determination with resolution=0.8. Cell types were assigned to each cluster in organoids with wild type by known gene markers and other organoids with knock-out gene were annotated according to wild type using function *MapQuery*.

The cell developmental trajectories were estimated using R package URD (v1.1.1)^54^. Firstly, the diffusion map was computed by function *calcDM* with KNN=100 and sigma.use=30. Next, based on RG cells, other cells were ordered in pseudotime using function *floodPseudotime* and *floodPseudotimeProcess* with the default parameters. The biased transition matrix was constructed using function *pseudotimeWeightTransitionMatrix* according to the difference in pseudotime between each pair of cells. Finally, the process of random walks was performed using *simulateRandomWalksFromTips* and the branching dendrogram structures were generated using function *bulidTree*.

### scRNA-seq differential expression analysis

Genes, which expressed in more than 10 cells, were used for further different expression analysis using the R package Seurat *FindAllMarkers*. Every comparison was performed between single KO and WT sample and genes with adjust p-value < 0.01 were defined as differentially expressed genes.

### Multi-electrode array (MEA) recording and analysis

The 12-week dorsal or ventral forebrain organoids were plated in 6-well MEA plates (Axion Biosystems) and each well contains 64 low-impedance platinum microelectrodes. Spontaneous neural spikes were recorded using the AxIS Navigator system and spikes were detected with an adaptive threshold crossing of > 6 standard deviation (s.d.) for each electrode. The organoids were first stayed for 5 minutes in the device, and then 3 minutes of data were recorded. We quantified the knock-out effects of gene on dorsal or ventral forebrain organoids based on MEA recording from three levels, including spike, electrode and network. Firstly, we used the empirical cumulative distributions to visualize the distribution of inter-spike intervals (ISIs) of different groups and then used the one-side Kolmogorov-Smirnov test to compare ISIs distributions of knock-out and wild-type group. Besides, we counted the firing rates of every electrode, *i.e.*, the number of spikes per minute. At neuron level, shorter ISIs and higher firing rates indicate stronger neuron activity. The synchrony index of network based on the whole organoid was used to estimate strength of neuron-neuron connections. The t-test was used to compare the knock-out effects of gene on firing rates of electrodes and synchrony of networks.

### *Caln1* knock-out mice

The C57Bl/6J Caln1^+/−^ mice, constructed by Cyagen Biosciences Company (Suzhou), were used as parent mice to generate offspring including WT, Caln1^+/−^ and Caln1^−/−^. To knock out mice Caln1, two gRNAs are used, *i.e.*, 5’-GCCATGGAGAGTGCTACTTA-3’ (GGG) and 5’-CATAGGTGACCCCGCCAAAG-3’(GGG). Three primers are used to verify knock-out mice, *i.e.*, 5’-TTAGAATATGCCAATGAAAGGGCTG-3’ (F1), 5’- ACCCCTATGACAAACACACATAAGA-3’ (R1) and 5’- GTTTGCTGCTGCTGTTATTATTGTG-3’ (R2).

### Western blotting

Protein samples were collected from WT or *Caln1^−/−^* mice cortex at the age of 14 weeks. A volume of 20 μl lysate with 40 μg protein was loaded for each sample. Protein samples were loaded in 15% SDS-PAGE and transferred to a PVDF membrane. The membranes were incubated in an orbital shaker at 4 ℃ overnight with the primary antibodies CALN1 (rabbit, 1:500, Proteintech, cat. #11477-1-AP), Cas9 (rabbit, 1:500, ThermoFisher, cat. #61978) and β-actin (rabbit, 1:2000, Cell Signaling Technology, cat. #8457). Horseradish peroxidase (HRP)-conjugated anti-rabbit IgG was used as a secondary antibody. Chemiluminescence (Yeasen, Super ECL Detection Reagent, cat. #36208ES60) was used to detect the signals.

### Mice neocortical slice preparation

Coronal slices of prefrontal cortex were obtained from adult WT C57/B6 mice and *Caln1^−/−^* mice (postnatal 9-16 weeks, male). The mice were anesthetized with pentobarbital sodium at a dose of 100 mg/kg by intraperitoneal injection, and then sacrificed with decapitation. The brain was then immediately dissected out and immersed in ice-cold oxygenated (95% O_2_ and 5% CO_2_) slicing solution in which 126 mM NaCl was substituted by 213 mM sucrose. The prefrontal cortical slices (250 μm in thickness) were cut with a microtome (VT-1200S, Leica, Germany) and incubated in the aerated normal ACSF (in mM): 126 NaCl, 2.5 KCl, 2 MgSO_4_, 2 CaCl_2_, 26 NaHCO_3_, 1.25 NaH_2_PO_4_, and 25 dextrose (315 mOsm, pH 7.4) and maintained at 34.5 °C for 1.5 hrs. After incubation, slices were kept in the same solution at room temperature until use.

### Electrophysiological recording

After slice preparation, the prefrontal cortical slices were transferred into a recording chamber and perfused with ACSF. The solution temperature was maintained at 34.5-35°C. Cortical neurons were visualized with an upright infrared differential interference contrast microscope (BX51WI; Olympus) equipped with a water-immersed objective (40×, numerical aperture 0.8). We performed whole-cell current-clamp recording from the soma of layer-V pyramidal cells to investigate their intrinsic membrane properties and excitability. The patch pipettes had an impedance of 5-6 MΩ when filled with a K^+^-based internal solution containing (mM) 140 K-gluconate: 3 KCl, 2 MgCl_2_, 10 Hepes, 0.2 ethylene glycol tetraacetic acid (EGTA), and 2 Na_2_ATP (290 mOsm, pH 7.23 with KOH). We also added 0.2% biocytin to the pipette solution for cell staining after the experiments. Electrical signals were acquired using a MultiClamp 700B amplifier (Molecular Devices), digitized and sampled by Micro 1401 mkII (Cambridge Electronic Design) at 25 kHz using Spike2 softwares.

### Analysis of electrophysiological and morphological data

An abrupt-burst spiking (ABS) event is defined as a sudden depolarization plateau with the generation of a burst of action potentials. We only counted ABS events occurred during the initial 1000-second recording. Spiking responses and the input-output (*I-F*) curves were examined by a series of positive current pulses (50 pA per step, 500 ms in duration). The threshold current (*i.e.*, rheobase) is defined as the current that only evokes a single action potential. The duration of individual ABS events is the period of depolarization above the baseline *V*m. Data was analyzed with MATLAB (MATHWORKS R2022b).

After electrophysical recording, slices were fixed with 4% paraformaldehyde for more than 24 hrs and then stained with Alexa Fluor-488 conjugated avidin. The z stack images (0.31 μm/pixel and 1 μm between successive images) of the neurons were acquired using a confocal microscope (A1/C2-DUVB, Nikon).

### Animal behavior test

Animal experiments were conducted according to the Guidelines for the Care and Use of Laboratory Animals of Fudan University and approved by Fudan University. C57Bl/6J mice were housed under the specific pathogen-free (SPF) conditions at a temperature of 22 ± 1℃ and a humidity of 55 ± 5%. Both sexes were used in behavior test and C57Bl/6J mice used as the social mice in the three-chamber social test were obtained from the Charles River Lab.

#### Behavioral measurements and video analysis

All behavioral analyses were performed in the Mouse Behavior Room of Institute for Translational Brain Research, Fudan University. Before each test, mice were acclimated within their housing cage in the behavior room for at least 30 min. The behavior cage was dimly illuminated at 30 lux or 100 lux due to different behavior tests. All the behavior test videos were analyzed by the ANY-maze software.

#### Y-maze test

The behavior cage include three arms (30 cm × 6 cm × 30 cm), with an angle of 120° between them. Arms were labeled as A, B and C and we defined arm A as the start arm, where all mice were placed in the distal part, facing back the center part of the maze. The tested mice were allowed to explore all three arms of the Y-maze for 10 min. The percentage of spontaneous alternation, total number of arm entries, total distance and average speed were calculated to measure the spatial working memory of mice. Spontaneous alternation (%) = Number of Alternations / (Total number of arm entries – 2) × 100.

#### Three-chamber social test

The three-chamber social apparatus is a rectangular white plastic box (60 cm × 40 cm × 22 cm), consisting of three equal chambers (left, middle and right chamber) with small doors in the walls between each chambers, and there were two small cages placed in diagonal inside the rectangular behavior box. During Phase 1, the habituation stage, subject mice were permitted to explore the middle part for 10 min under the condition of both side chamber doors closed. At Phase 2, one stranger mouse was placed into one of the cages in diagonal, and the other cage was still empty. The doors to the left and right chambers were raised and tested mouse was allowed to explore all three chambers for 10 min (Session 1). The subject mouse was then briefly expelled to the middle chamber while another new stranger mouse was placed in the other previously empty cage. At Phase 3, doors were raised, and the subject mouse was allowed to explore all three chambers for 10 min (Session 2). The sex, strain and age of stranger mice were the same as our subject mice. The ANY-maze software was used to record the activity trajectory of tested mice and the time of some important events including shuttling between chambers and effective social activities with other mouse, which is, the nose of tested mouse facing towards stranger mouse within 20 mm. We analyzed the social duration in session 1 and 2 and duration in middle area to assess the social preference of KO mice.

#### Open-field test

The cage with a square base (40 cm × 40 cm) and four 40-cm high walls was used in the open-field test. The tested mice were placed in the middle of the cage, and were allowed for free exploration in the environment for 10 min. The activity was recorded by camera in behavior cage and analyzed by ANY-maze software. The percentage of duration and total moving distance in center area were used to analyze the level of anxiety and locomotion.

#### Elevated plus maze test

The apparatus located 1 meter high above the floor is made up of two open arms (25 × 5 × 0.5 cm) and two closed arms (25 × 5 × 16 cm) across from each other at the center. The behavior room is soundproof and the illumination level is maintained at 100 lux. The tested mice were softly placed in the center area of the maze with their head facing towards the open arm and were allowed to move freely for 10 min. The activity was recorded by a camera above the maze. Percentage of duration in the open arm was calculated to provide measurements of anxiety-like behavior.

#### Tail suspension test

This test is used to create a despair state to induce the desire of living and to test level of depression in mice. Mice were suspended by the tail with tapes above a flat surface in the tail suspension cage (BIO-TST5, BIO-SEB). After the habituation stage (*i.e.*, 1 min), by the software BIO-TST5 and BIO-SEB, behavioral scores of mice were obtained basing on the duration of immobility within 5 min.

#### Pre-pulse inhibition (PPI) test

The whole process of PPI was monitored using SR-Lab chambers. Mice were acclimated in the PPI apparatus for 5 min in the presence of a 65-dB background noise and then the test session was automatically started. Each test session began with 10 trials of the startle stimulus (*i.e.*, 120 dB) alone, followed by 30 combinatorial trials of a pre-pulse stimulus (*i.e.*, 72, 76 and 80 dB) along with a startle stimulus, of which three intensities of pre-pulse stimulus were conducted randomly and the total number of each pre-pulse stimulus was 10. The intertrial intervals (ITIs) varied from 8 to 20 sec. In the PPI stage, each pre-pulse and startle stimulus lasted 20 ms and 40 ms, respectively, and the intervals between pre-pulse stimulus and startle stimulus were 80 ms. Data of transient response from tested mice were collected after each startle stimulus. The percentage of PPI = (1 –PPi_S/S) × 100, where PPi_S indicates the pre-pulse intensity along with a startle stimulus, S indicates the startle-only trials.

#### SEP-363856 treatment test

*Caln1^−/−^* mice were administrated with SEP-363856 (MCE, cat. # HY-136109, 10mg/kg) via oral gavage once a day for 3 days. Behavior videos were collected and analyzed before and after each oral gavage administration.

## Acknowledgments

This study was supported by the STI2030-Major Projects (Z.S., Y.S., 2021ZD0202500), National Natural Science Foundation of China (Z.S., 32070956; Y.S., 32130044, T2241002; HJ.L., 82101574; X.L., 32200807), China Postdoctoral Science Foundation (HJ.L., 2021TQ0073, 2021M700837), Program of Shanghai Academic/Technology Research Leader (Y.S., 21XD1400100), MOE Frontiers Center for Brain Science fund and starting fund from Fudan Universities (Z.S.).

## Authors contributions

Z.S. and HJ.L. designed the experiments. HJ.L., T.C. and J.C. performed sgRNAs design and CRISPR-Cas9 plasmid construction. HJ.L. constructed knock-out cell lines. HJ.L. induced organoids. HJ.L. and J.X. performed scRNA-seq experiments. HJ.L. performed bulk RNA-seq and scRNA-seq analysis. HJ.L. performed organoids immunostaining. HJ.L. performed MEA experiments and analysis. X.Y. performed animal experiments and analysis. X.L. performed animal electrophysiological studies, supervised by Y.S. Z.S., HJ.L., X.Y., X.L., S.C. and Y.S. wrote the manuscript and data interpretation. Z.S supported this study financially.

## Data and materials availability

All data are available in the main text or the supplementary materials. All sequence data in this study have been deposited within the Genome Sequence Archive for Human (https://ngdc.cncb.ac.cn/gsa-human/) with the accession identifier HRA005215.

## Code availability

The present study did not use customized mathematical algorithm and the parameters of software used during the analysis have been described in detail in the Methods section.

---

**Figure S1.**
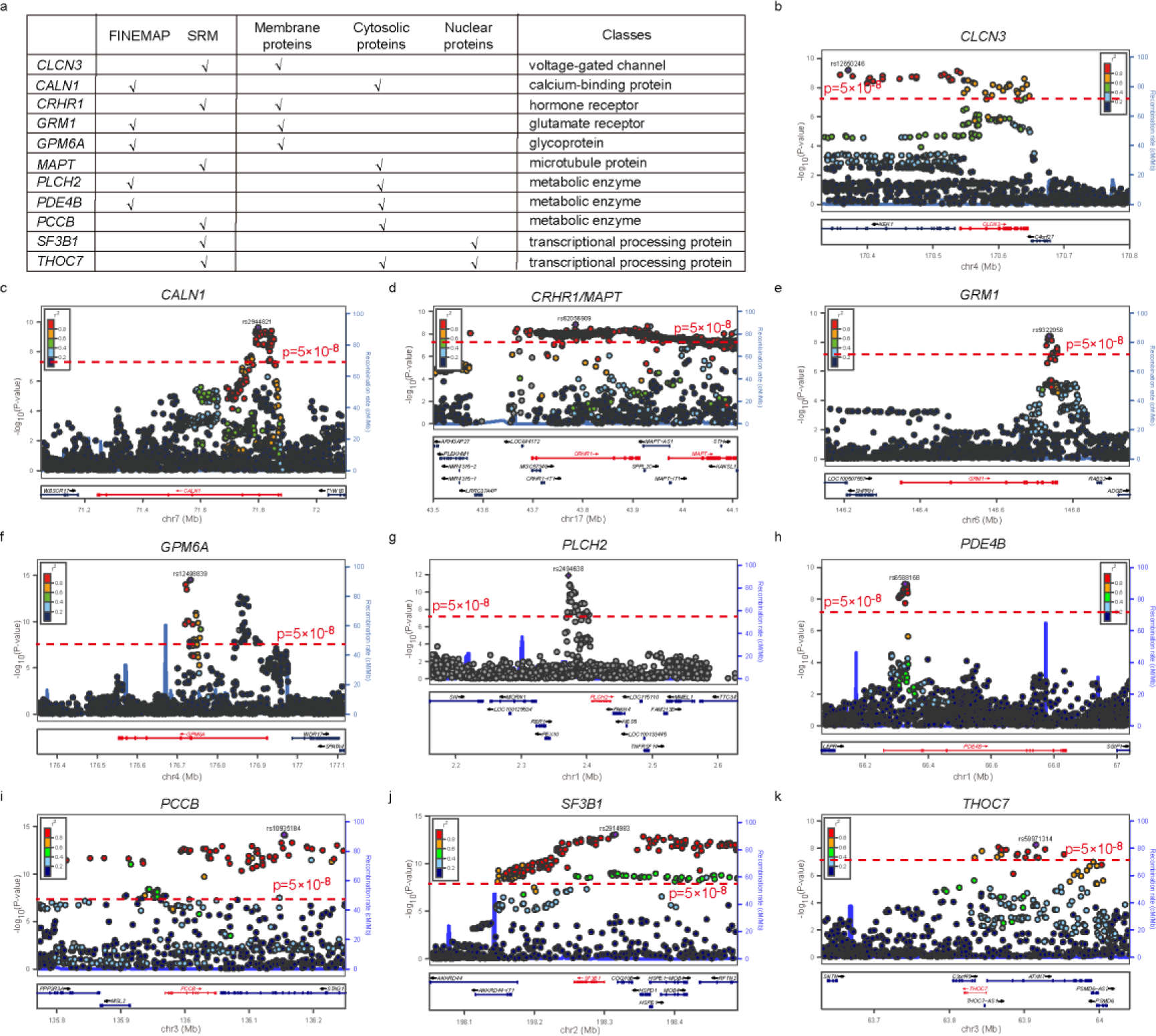
The classes and associations with disease of 11 selected schizophrenia risk genes. **a**, The classes of risk genes. **b-k**, The regional association plots of risk genes from the latest schizophrenia GWAS which based on 74,776 cases and 101,023 controls from European, East Asian, African American and Latino ancestry.

**Figure S2.**
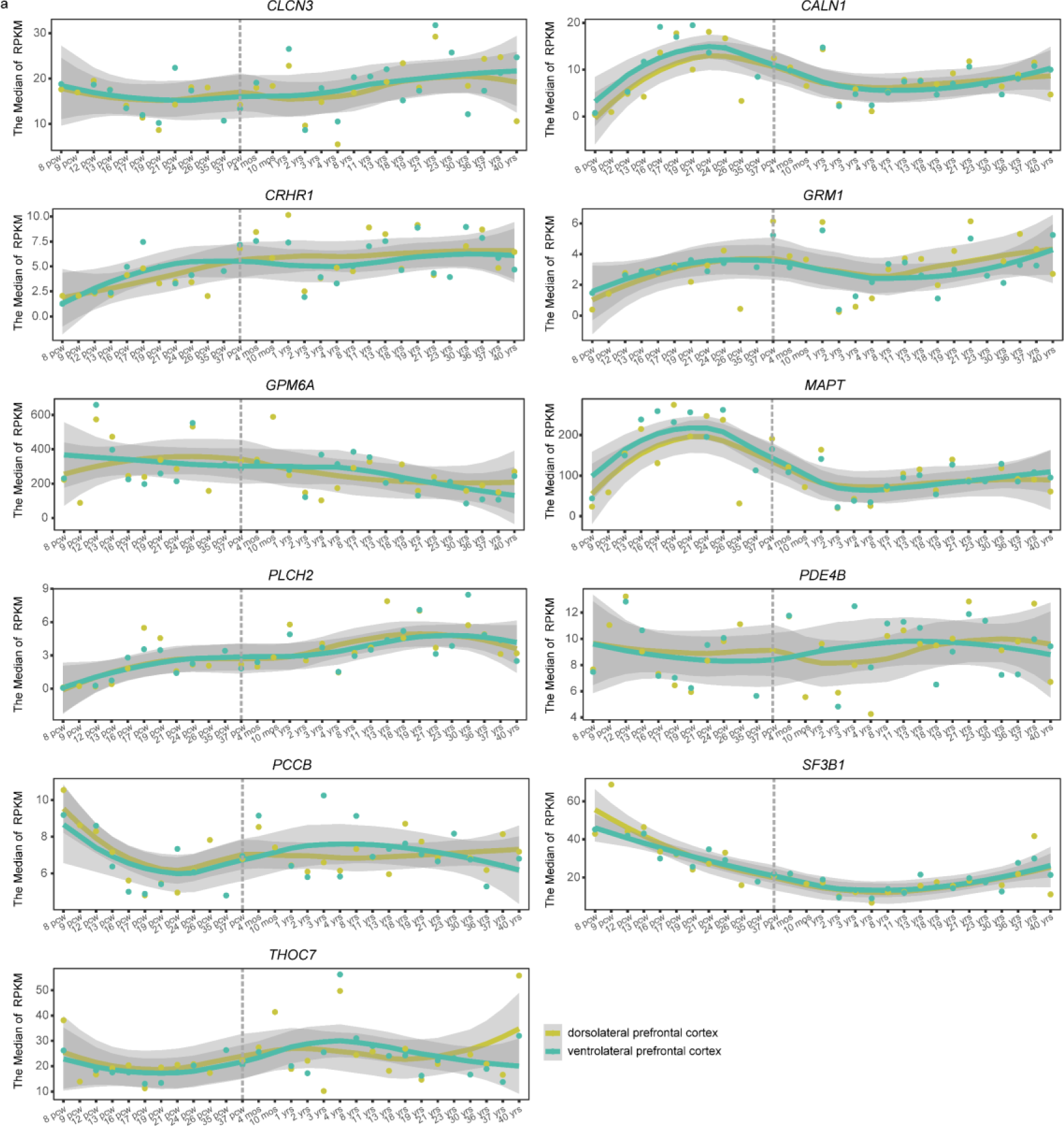
The human dorsolateral and ventrolateral prefrontal cortex expression trajectory of 11 selected schizophrenia risk genes across development stages. **a**, Based on the BrainSpan database, the smooth lines of gene expression RPKM values from human 8 pcw to 40 yrs were displayed. The dotted line indicates before and after human birth. RPKM, Reads Per Kilobase per Million mapped reads. pcw, post-conception weeks. mos, months. yrs, years old.

**Figure S3.**
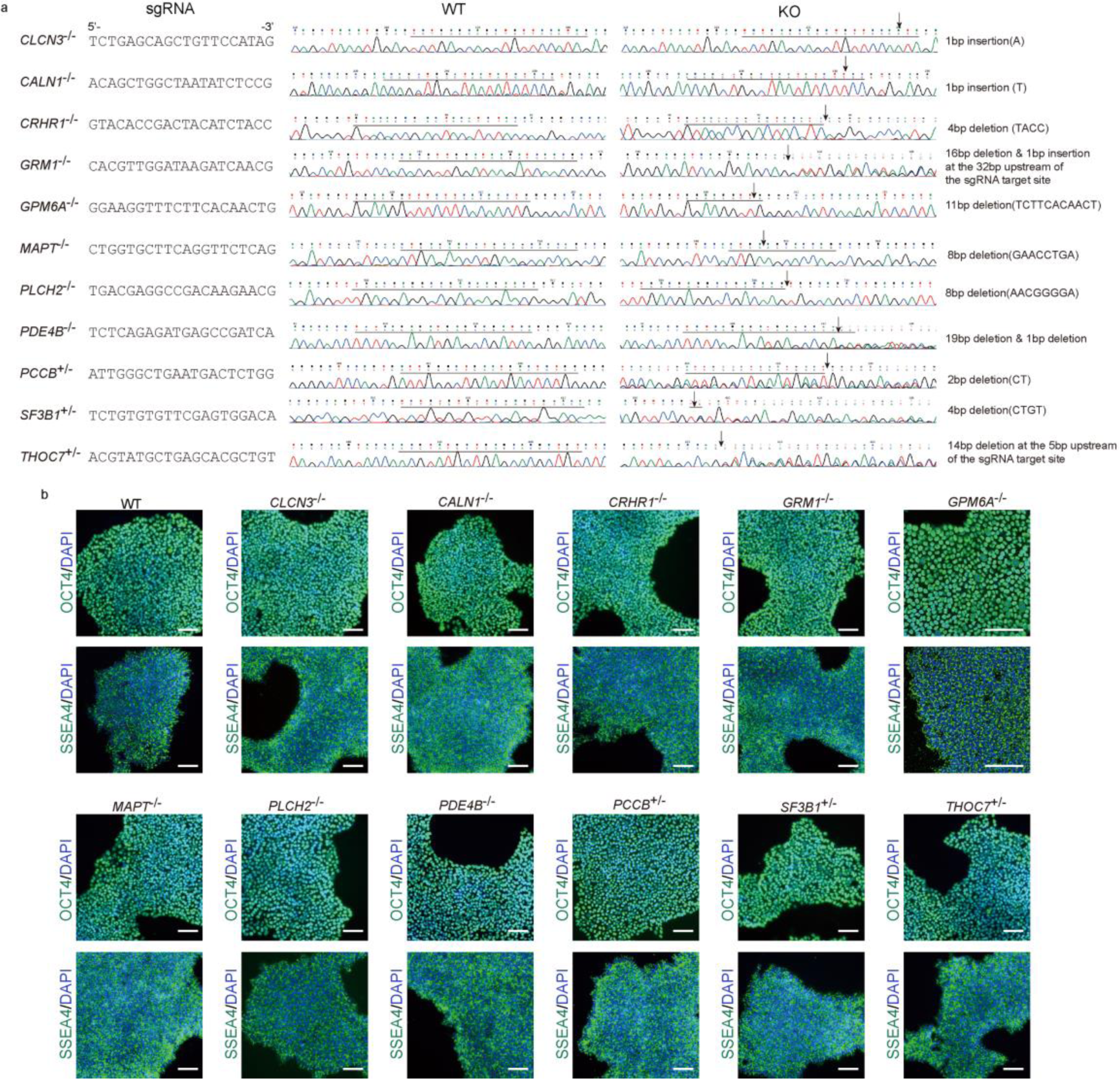
Sanger sequence and validation of pluripotency of knock-out hESC lines. **a**, Mutant type of each KO hESC line was verified by Sanger sequence. **b**, Immunostaining of OCT4 and SSEA4 in each KO hESC line used in this study.

**Figure S4.**
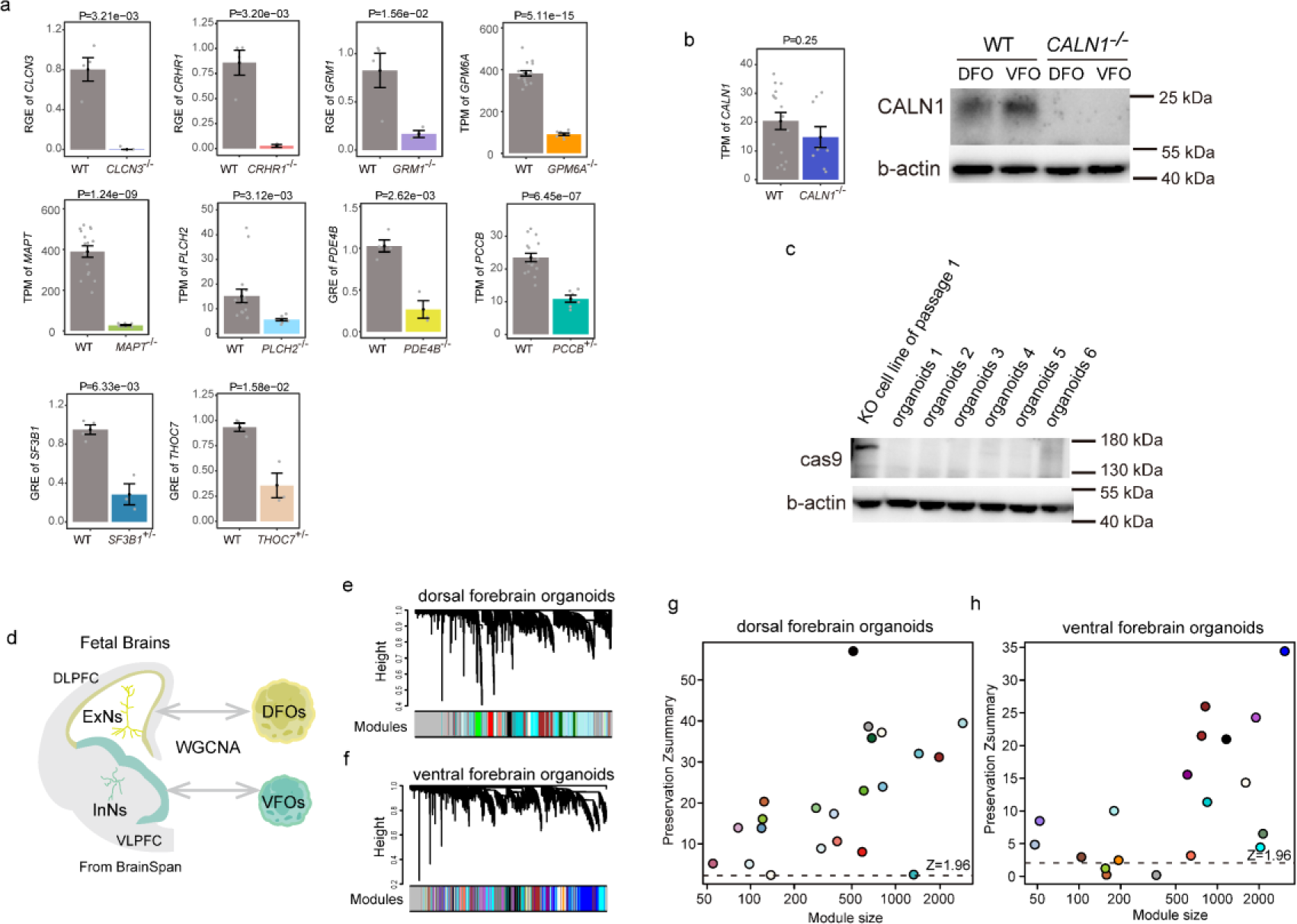
Expression of target gene in KO organoids and weight gene co-expression network analysis (WGCNA) in 6-week DFOs and VFOs. **a,** Normalized gene expression values of target gene in KO 6-week-old organoids. TPM (from RNA-seq), transcripts per million. RGE (from qPCR), relative gene expression. **b,** CALN1 loss at protein level in *CALN1^−/−^* DFOs and VFOs. **c,** The Cas9 is silencing in knock-out organoids. **d**, Comparison between forebrain organoids and fetal prefrontal cortex based on RNA-seq data. DLPFC, dorsolateral prefrontal cortex. VLPFC, ventrolateral prefrontal cortex. WGCNA, weight gene co-expression network analysis. **e, f**, A total of 22 and 19 gene expression modules (except for the gray module) were identified by WGCNA in dorsal (**e**) and ventral (**f**) forebrain organoids, respectively. **g, h**, The Z-summary statistics of each module in the dorsal (**g**) and ventral (**h**) organoids comparing with DLPFC and VLPFC of fetal brains. The modules with Z-score > 1.96 were considered as preservative.

**Figure S5.**
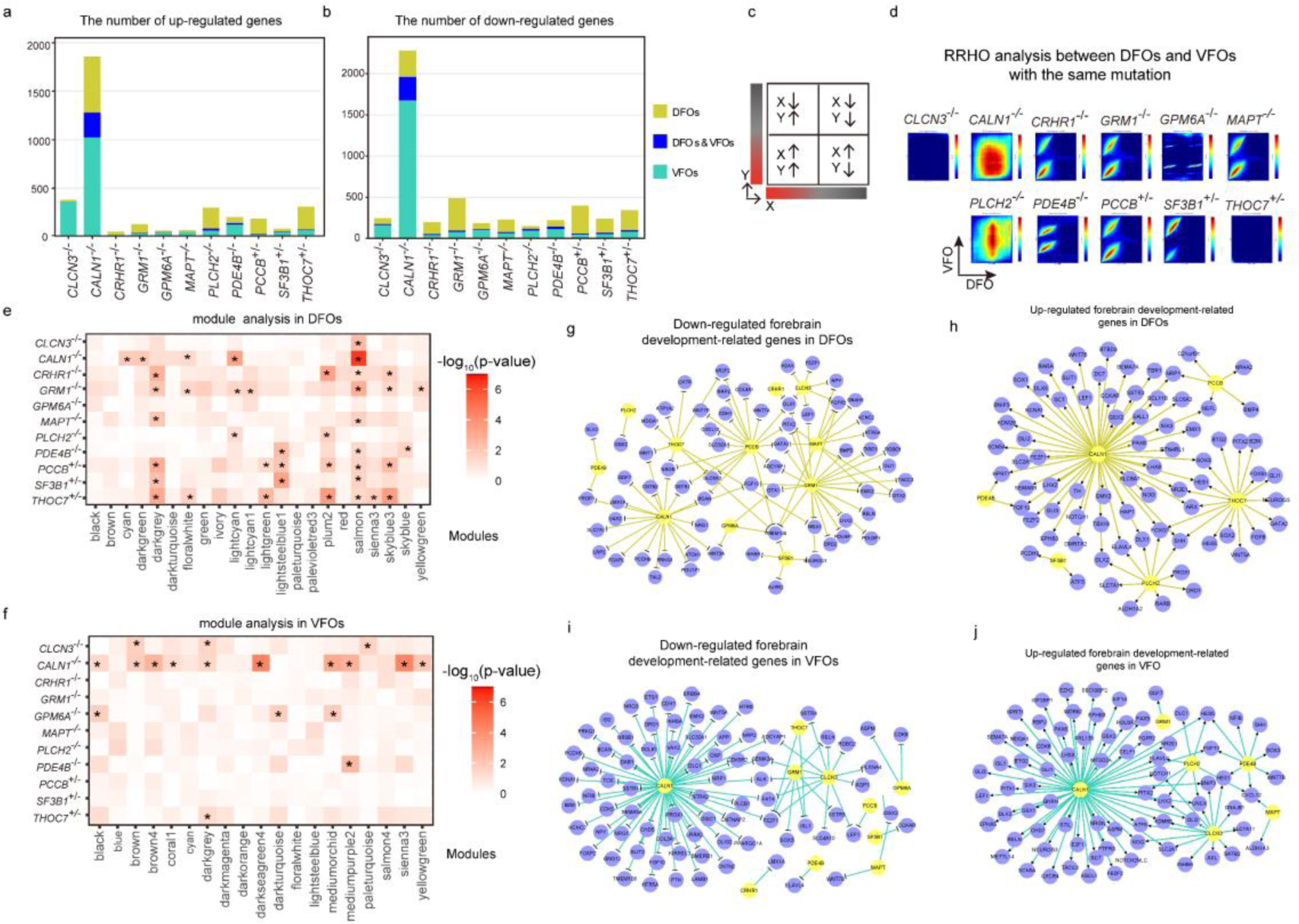
Bulk RNA-seq analysis of different schizophrenia risk genes in dorsal and ventral forebrain organoids at 6 weeks. **a, b**, The statistics of up-regulated (**a**) and down-regulated (**b**) genes for each KO gene. DFOs & VFOs, common up- or down-regulated in dorsal and ventral forebrain organoids. **c**, Interpretation of RRHO map. **d**, The RRHO maps comparing between DFOs and VFOs with the same KO gene. Signals in top right quadrant and bottom left quadrant indicate an overlap of down-regulated and up-regulated genes found in both DFOs and VFOs, respectively. **e, f**, Knockout of schizophrenia risk genes impact different gene co-expression modules in DFOs (**e**) and VFOs (**f**). The p-values were calculated using two tailed t-test. *, p-value < 0.05. **g-j**, The Differentially expressed genes related with forebrain development in KO DFOs (**g, h**) and VFOs (**i, j**).

**Figure S6.**
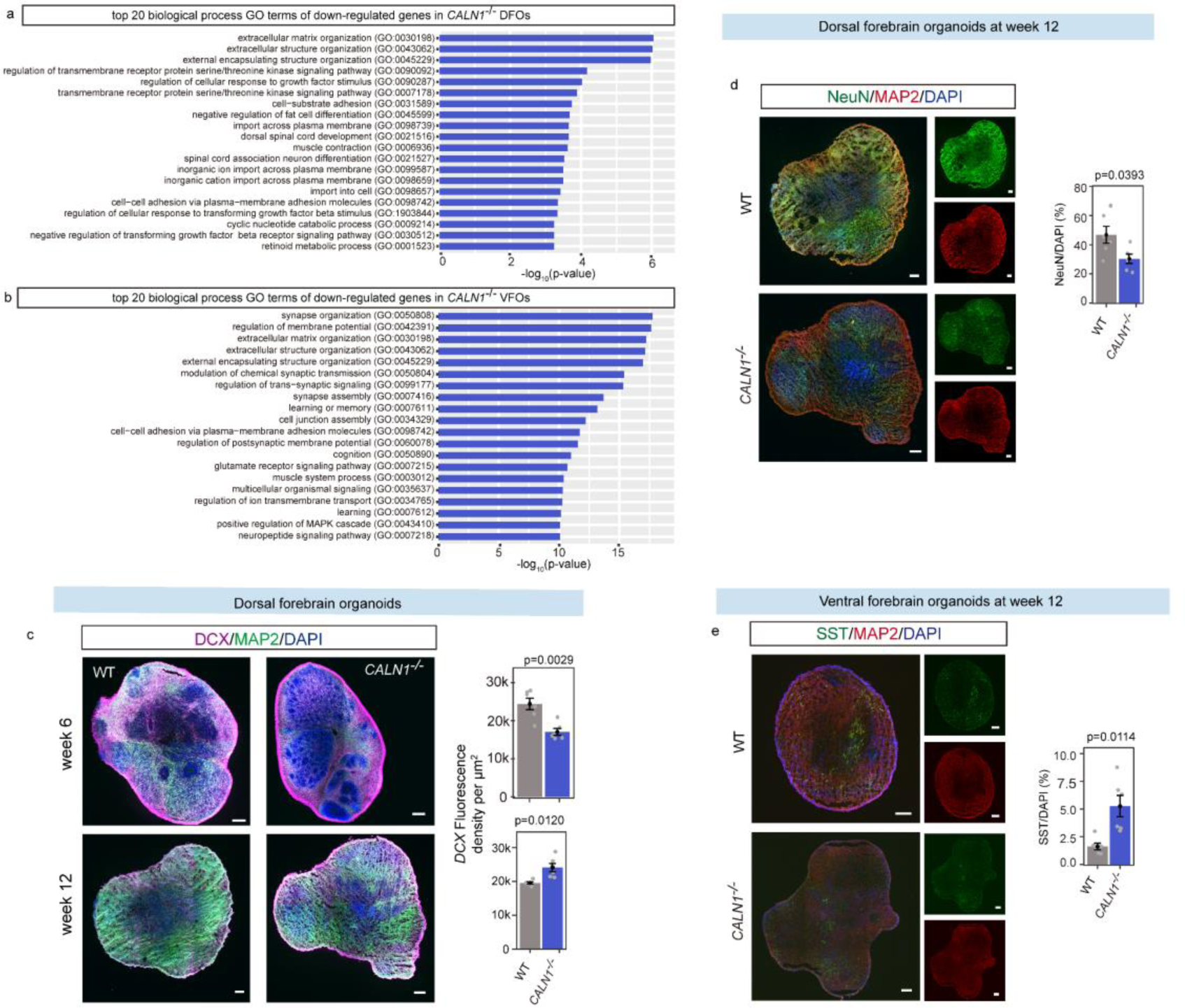
Knockout of *CALN1* impacts specific GO terms and cell types. **a, b,** Top 20 GO terms related with biological process enriched by down-regulated genes in *CALN1^−/−^* DFOs (**a**) and VFOs (**b**). **c-f,** Immunostaining of specific cell markers, including DCX (**c**) and NeuN (**d**) in *CALN1^−/−^* DFOs and SST (**e**) in *CALN1^−/−^*VFOs. The p-values were calculated using two tailed t-test. scale bar, 200 μm.

**Figure S7.**
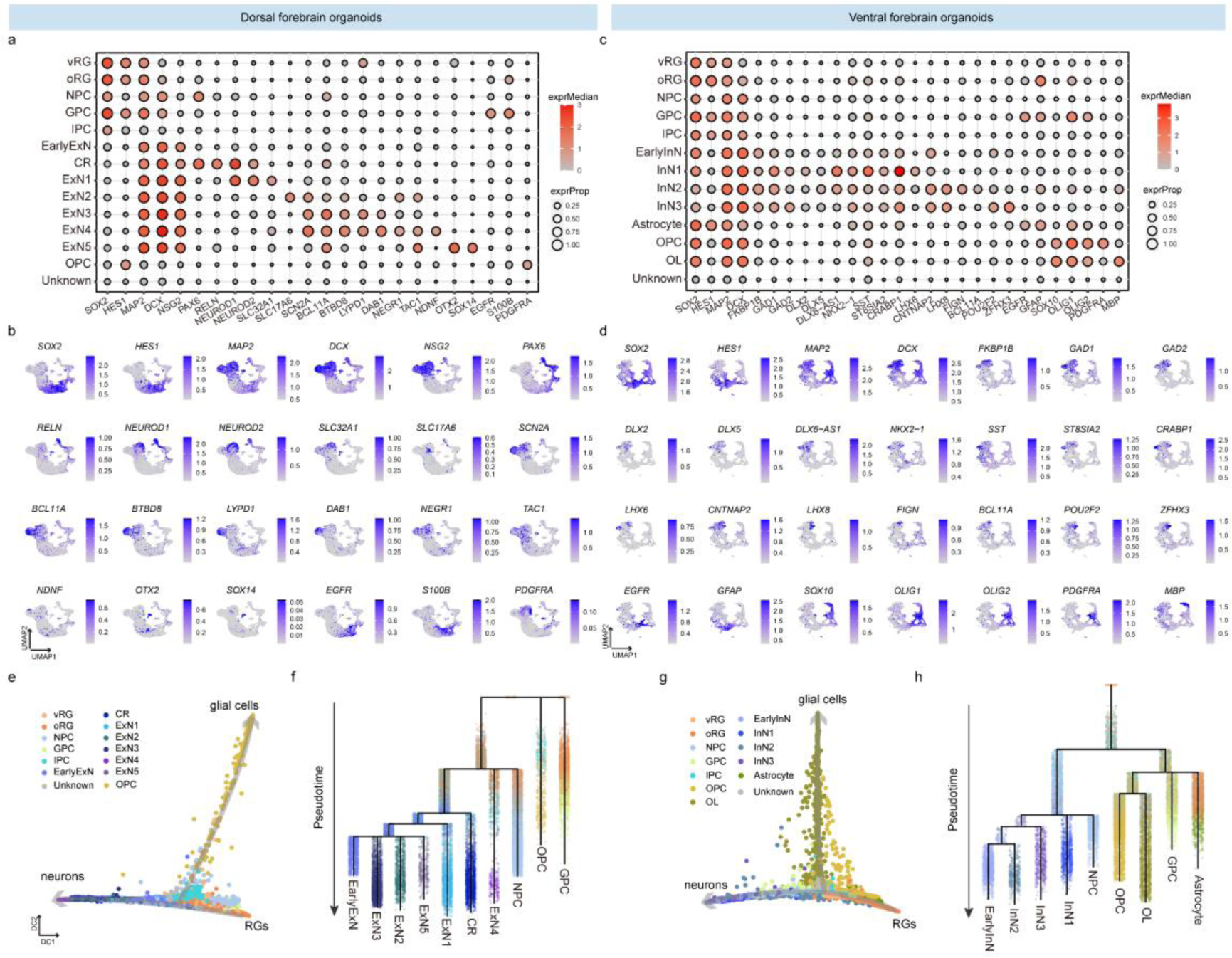
Characteristics of cell types and development trajectories. **a, c,** Annotation of cell clusters according to known marker genes for DFOs (**a**) and VFOs (**c**). **b, d**, The UMAP plots of known marker genes for DFOs (**b**) and VFOs (**d**). **e, g**, The diffusion maps of different cell clusters for DFOs (**e**) and VFOs (**g**). RGs, radial glial cells. **f, h**, The branching dendrogram structures of neurons and glial cells for DFOs (**f**) and VFOs (**h**).

**Figure S8.**
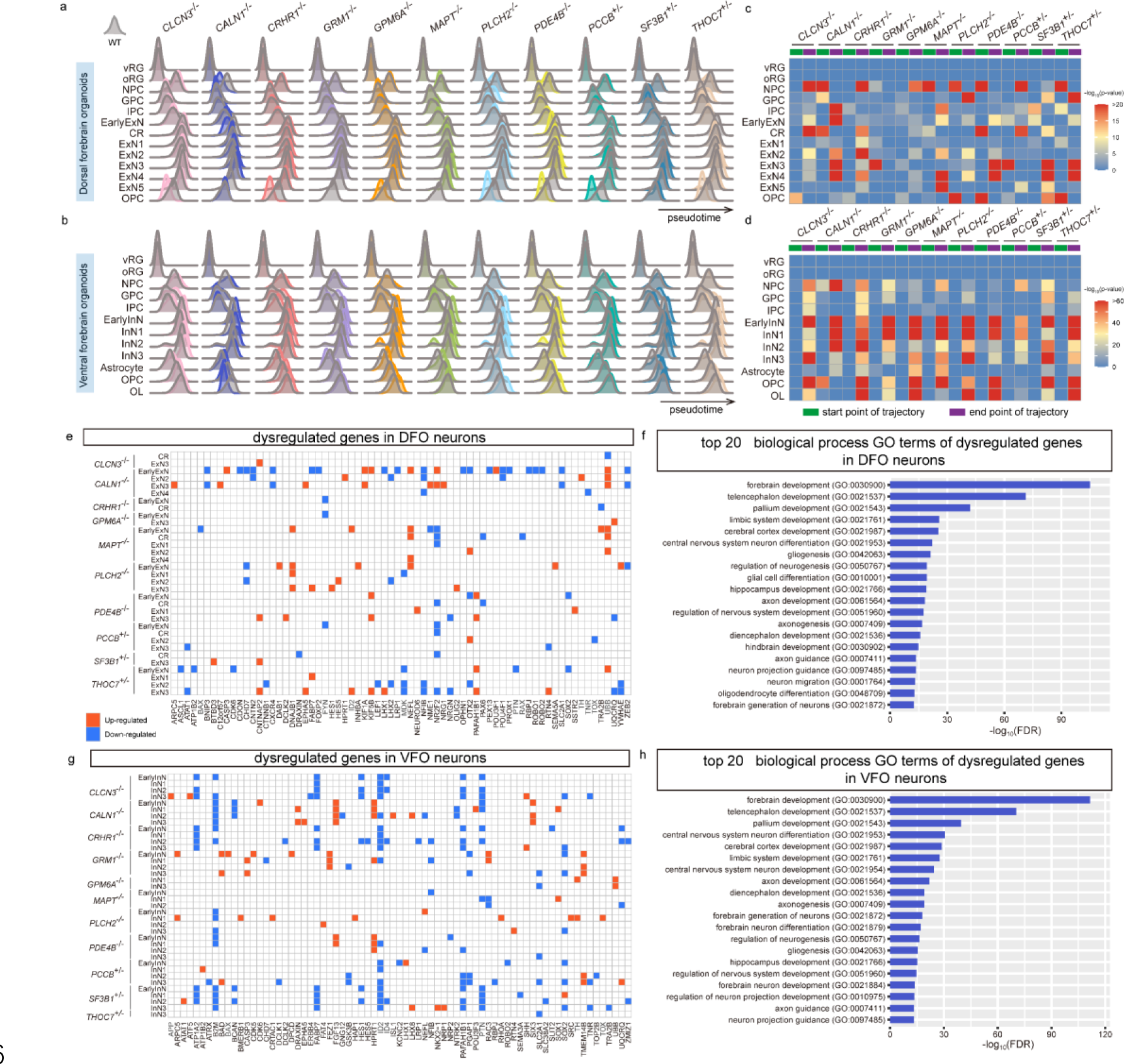
Schizophrenia risk genes affect organoids development trajectories and gene expression of neurons. **a, b**, The density plots of pseudotime for each cell cluster in KO DFOs (**a**) and VFOs (**b**). **c, d**, Compared with wild type, the maturation of cell clusters in KO DFOs (**c**) and VFOs (**d**) were estimated using one-tailed Kolmogorov-Smirnov tests. The increasing tendency to the start and end point of trajectory indicate delayed and accelerated maturation, respectively. **e, g**, Dysregulated genes in specific neuron clusters in KO DFOs (**e**) and VFOs (**g**). **f, h**, Top 20 GO terms related with biological process enriched by dysregulated genes in DFO neurons (**f**) and VFO neurons (**h**).

**Figure S9.**
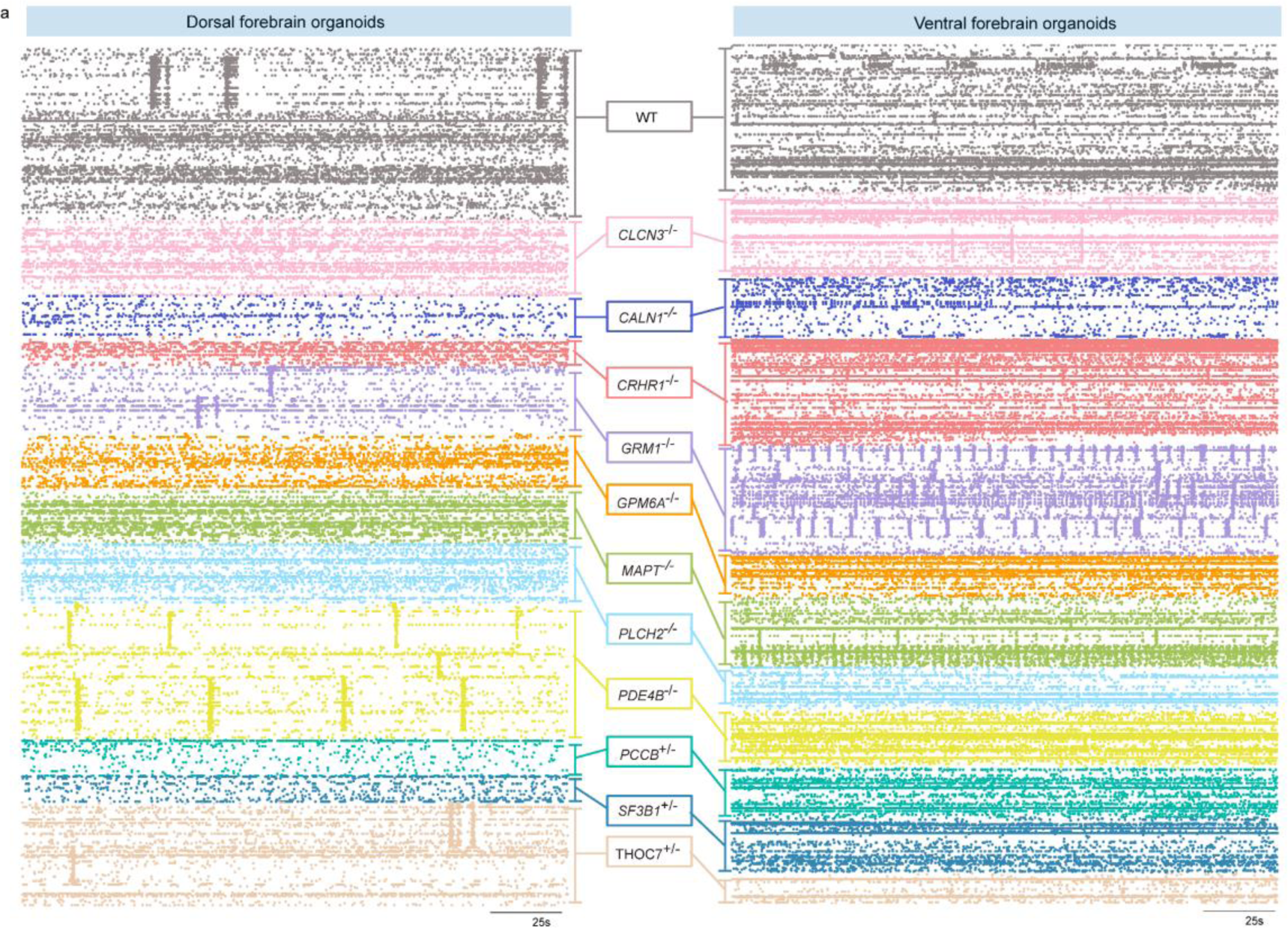
Spike spots of MEA across individually active electrode. **a,** A total of 61,654 and 88,850 spikes were generated by 962 and 893 active electrodes in DFOs and VFOs, respectively.

**Figure S10.**
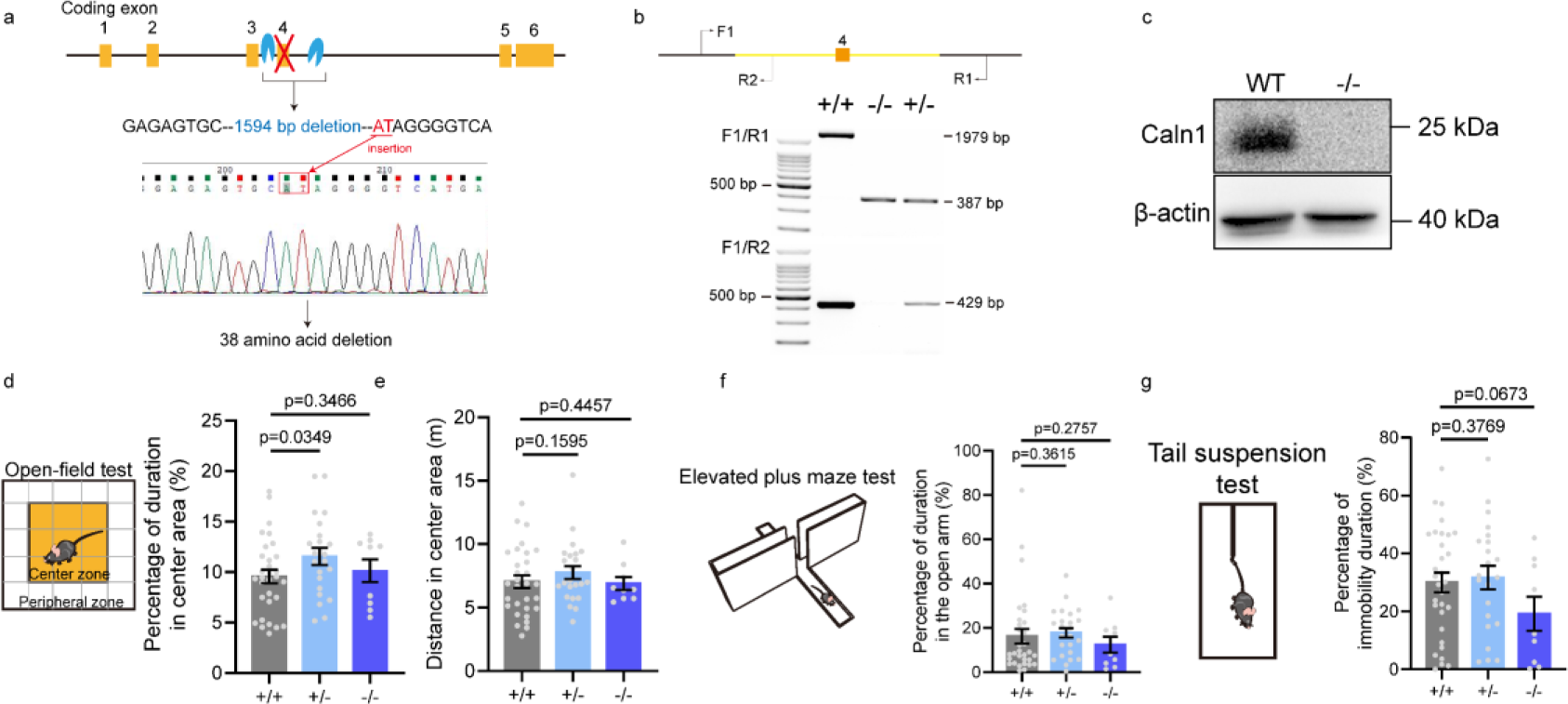
The validation of *Caln1^−/−^* mice and anxiety and depression behavior tests. **a-c**, *Caln1* knock-out mice were verified by Sanger sequence (**a**), DNA agarose gel electrophoresis (**b**) and Western blot (**c**)**. d**, **e**, The open-field test**. f**, The elevated plus maze test. **g**, The tail suspension test. Data are presented as mean ± SE. The one-tailed t-test was used to calculate p-value.

**Figure S11.**
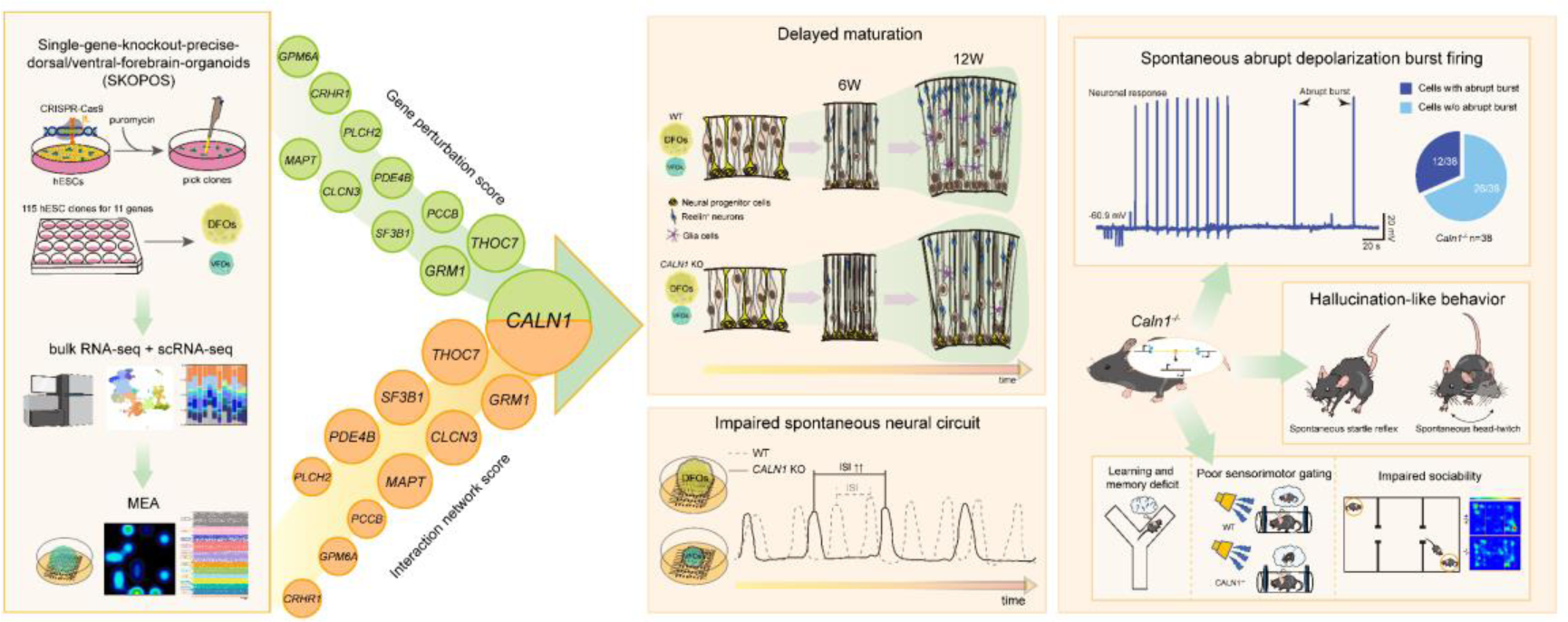
Conceptual schematic diagram underlining the main results in this study. Calneuron 1 (*CALN1*), screened by single-gene-knockout-precise-dorsal/ventral-forebrain-organoids (SKOPOS) system, reveals the pivotal roles in schizophrenia via perturbing human forebrain organoids development and causing hallucination-like behavior in mice.

